# IgEvolution: clonal analysis of antibody repertoires

**DOI:** 10.1101/725424

**Authors:** Yana Safonova, Pavel A. Pevzner

## Abstract

Constructing antibody repertoires is an important error-correcting step in analyzing immunosequencing datasets that is important for reconstructing evolutionary (clonal) development of antibodies. However, the state-of-the-art repertoire construction tools typically miss low-abundance antibodies that often represent internal nodes in clonal trees and are crucially important for clonal tree reconstruction. Thus, although repertoire construction is a prerequisite for follow up clonal tree reconstruction, the existing repertoire reconstruction algorithms are not well suited for this task. Since clonal analysis has the potential to reveal errors in the constructed repertoires and contribute to constructing more accurate repertoires, we advocate a *tree-guided construction of antibody repertoires* that combines error correction and clonal reconstruction as interconnected (rather than independent) tasks. We developed the IgEvolution algorithm for simultaneous repertoire and clonal tree reconstruction and applied it for analyzing multiple immunosequencing datasets representing antigen-specific immune responses. We demonstrate that analysis of clonal trees reveals highly mutable positions that correlate with antigen-binding sites and light-chain contacts in crystallized antibody-antigen complexes. We further demonstrate that this analysis leads to a new approach for identifying complementarity determining regions (CDRs) in antibodies.

## Introduction

Diversification of antibody repertoires is achieved by various processes that include *V(D)J recombination*, *intergenic insertions*, *heavy and light chain pairing*, and *somatic hypermutagenesis*. This paper focuses on somatic hypermutagenesis that introduces *somatic hypermutations* (*SHMs*) into evolving antigen-specific antibodies (*secondary diversification* of antibodies). After a successful antibody-antigen binding, the corresponding B-cell undergoes a cell division in a lymph node (*clonal expansion*) that aims to increase the pool of antibodies that bind to an antigen. At the same time, the *activation-induced deaminase* is activated in such B-cell and its clones to introduce SHMs into immunoglobulin genes. Although SHMs change the three-dimensional structure of an antibody (and thus its ability to bind to an antigen) in a random way, the regulatory mechanisms of the immune system play the role of the natural selection for maximizing the antigen-binding abilities of antibodies. They expand B-cells with high affinity to an antigen and kill *self-reactive* B-cells with potentially harmful mutations (Figure S1).

Secondary diversification turns a naive antibody repertoire into a set of *clonal lineages*, where each clonal lineage is formed by descendants of a single naive B-cell. The immunoglobulin genes within the same clonal lineage share the same V(D)J recombination (including intergenic insertions and exonuclease removals) and differ by SHMs only. Reconstructing evolution of antibody repertoires using immunosequencing (*Rep-Seq*) datasets has important biomedical applications, such as evaluating vaccine efficiency, monitoring immune response, and optimizing antibody drugs.

Evolution of B-cells in each clonal lineage is described by a *clonal tree*, where each vertex corresponds to a B-cell and each B-cell is connected by a directed edge with all its immediate descendants. Antibodies evolve under selection pressure that often results in *convergent evolution* (Thorsélius et al., 2006; Parameswaran et al., 2013; Jackson et al., 2014) when similar or identical antibodies are present in different repertoires. In the case of a single repertoire, convergent evolution is manifested in the identical mutations at the same position that occur multiple times in different branches of the clonal tree. Moreover, different paths in the tree may result in the same antibody produced by different B cells (Krause et al., 2011; Wu et al., 2011). Analysis of convergent evolution within a single clonal lineage remains an open computational problem that requires accurate algorithms for clonal tree reconstruction.

Emergence of Rep-seq technologies triggered developments of various clonal tree reconstruction approaches (Barak et al., 2008, Davidsen and Matsen, 2018) such as likelihood-based ImmuniTree (Sok et al., 2013) and IgPhyML (Hoehn et al., 2017) algorithms. Horns et al., 2016 proposed a *minimum spanning tree* (*MST*) algorithm for clonal tree reconstruction and applied it to barcoded Rep-seq datasets that were error-corrected using unique molecular identifiers (a similar MST-based BRILIA tool was developed by Lee et al., 2017).

These approaches, while useful, have not adequately addressed the challenge of dealing with sequencing and amplification errors in Rep-seq datasets. Some algorithms for clonal tree reconstruction (such as IgPhyML) even assume that the antibody repertoire is error-free and thus propagate errors in Rep-seq datasets into the constructed clonal trees. Since various sample preparation artifacts introduce artificial diversity in Rep-seq datasets (thus complicating follow-up clonal tree reconstruction), correcting errors in these datasets (repertoire construction) is a prerequisite for all clonal tree reconstruction algorithms. However, Shlemov et al., 2017 demonstrated that the state-of-the-art repertoire construction tools perform well on *background* repertoires (that have not undergone the selection process after stimulation by the antibody-antigen binding) but deteriorate on *stimulated* repertoires (e.g., repertoires evolved in response to vaccination or disease), arguably the most important type of repertoires for various biomedical studies. Although some Rep-seq protocols include molecular barcoding step, most publicly available Rep-seq datasets (including studies of stimulated repertoires) do not contain molecular barcodes and thus remain poorly analyzed from the point of view of clonal tree reconstruction.

We thus argue that generation of an antibody repertoire and clonal tree reconstruction are two interconnected problems that should be addressed together rather than independently as before. To integrate these problems, we developed the IgEvolution algorithm for simultaneous error correction and clonal tree reconstruction. Extending the BRILIA algorithm (Lee et al., 2017), IgEvolution decomposes a Rep-seq dataset into clonal lineages, constructs an MST for each clonal lineage, and analyzes leaves in the constructed trees to identify erroneous sequences. In difference from BRILIA that arbitrarily selects one of many MSTs (and does not correct some erroneous sequences), IgEvolution finds an MST that maximizes the number of leaves (leading to a more comprehensive error correction), and further performs an *iterative* (rather than a single round as in BRILIA) error correction.

We applied IgEvolution for constructing clonal trees from multiple immunosequencing datasets. To analyze the constructed clonal trees, we introduced the concept of *clonal mutability* of each position in an antibody and analyzed whether highly mutable positions correlate with the antigen-binding sites and light-chain contacts in crystallized antibody-antigen complexes. Since paired datasets that contain information about an antibody repertoire and a crystallized antibody-antigen complex resulting from this repertoire are currently not available, we analyzed antigen-binding sites and light-chain contacts in all publicly available crystallized antibody-antigen complexes. Even though our Rep-seq datasets originate from different sources than antibody-antigen complexes, we show that highly mutable positions and positions with convergent SHMs derived from the clonal trees correlate with antigen-binding sites and light-chain contacts.

We further use the concept of mutability to refine the bounds of CDRs computed using the existing similarity-based methods (Kabat et al., 1979; Chothia and Lesk, 1987; Giudicelli et al., 1997). We demonstrate that CDRs identified by these algorithms often miss some antigen-binding sites, leading to “shrinkage” of these important regions. Also, these algorithms do not reflect the fact that some regions of CDRs rarely contribute to antigen-binding sites as compared to other regions, e.g., the first half of CDR1 contributes to much fewer antigen-binding sites than its second path. Analyzing highly mutable positions in clonal trees provides an alternative to the classical similarity-based approach for identifying CDRs.

Finally, we illustrate how analysis of clonal trees may contribute to revealing alleles of germline IGHV genes that are associated with specific immune responses.

## Methods

### Limitations of existing repertoire construction tools

Repertoire construction is an important step for correcting sample preparation errors and finding putative receptor sequences among error-prone sequences in Rep-seq datasets. The existing repertoire construction tools, such as pRESTO (Vander Heiden et al., 2014), MiXCR (Bolotin et al., 2015), and IgReC (Shlemov et al., 2017) define the *abundance* of an antibody as the number of reads with the same (or nearly the same) sequence and further report high-abundance antibodies as putative receptor sequences. As a result, they typically reconstruct sequences from abundant B-cells but often miss low-abundance sequences representing evolutionary development of these B-cells that are important for clonal tree reconstruction.

In the past, clonal analysis used the constructed repertoire to infer the clonal trees but never changed the initially constructed repertoire based on the insights gained during the clonal analysis. This is unfortunate since clonal analysis has the potential to reveal errors in the constructed repertoires and contribute to constructing more accurate repertoires. A more accurate repertoire allows one to generate a more accurate clonal tree, thus resulting in a feedback loop between repertoire construction and clonal analysis. We thus advocate a *tree-guided construction of antibody repertoires* that combines error correction and clonal reconstruction of an antibody repertoire as interconnected (rather than independent) tasks.

### Hamming graphs

Safonova et al., 2015 introduced the concept of the *Hamming graph (HG)* for representing similarities between sequences in a Rep-seq library. All reads with the same sequence in a library corresponds to a vertex in the HG and two vertices are connected by an (undirected) edge if the Hamming distance between them does not exceed a *distance threshold*. The *weight* of an edge is defined as the Hamming distance between vertices it connects. The *abundance* of a vertex is defined as the number of reads that contributed to this vertex.

Since repertoire reconstruction tools attempt to identify reads derived from *identical* immunoglobulins, the IgReC tool (Safonova et al., 2015; Shlemov et al., 2017) uses a rather small default distance threshold for defining edges in the HG. Since clonal reconstruction tools attempt to identify reads derived from *clonally related* (rather than identical) immunoglobulins, IgEvolution increases this threshold as described in Supplemental Note “Constructing Hamming graphs for follow-up clonal tree reconstruction”.

### Tree-guided construction of antibody repertoires

IgEvolution constructs an MST of the HG and further analyzes it with the goal to iteratively remove vertices representing amplification and sequencing errors. A vertex in an MST is classified as a *low-abundance vertex* if either (i) its abundance is below *minAbundance* threshold (the default value is 5) or (ii) its abundance is at least *ratioAbundance* times lower than the abundance of one of its neighbors (the default value is 20). Supplemental Note “Setting a threshold for detecting low-abundance receptor sequences” describes how IgEvolution automatically selects a dataset-specific abundance thresholds depending on the number of PCR cycles applied during the sample preparation.

Analysis of various immunosequencing datasets revealed that the vast majority of low-abundance vertices represent leaves of the MST. Moreover, many low-abundance leaves are organized into large *star subgraphs*, i.e., are connected to the same vertex in an MST. Figure 1 shows an MST of the largest HG for the of FLU1-4 dataset that contains 125,050 reads sampling a heavy chain repertoire taken from the same individual at different time points after flu vaccination (donor 4 from the FLU1 dataset described in Table S2). IgEvolution constructed 2406 clonal lineages, with the largest clonal lineage containing 12,445 (328) reads before (after) removing low-abundance leaves.

**Figure 1.**
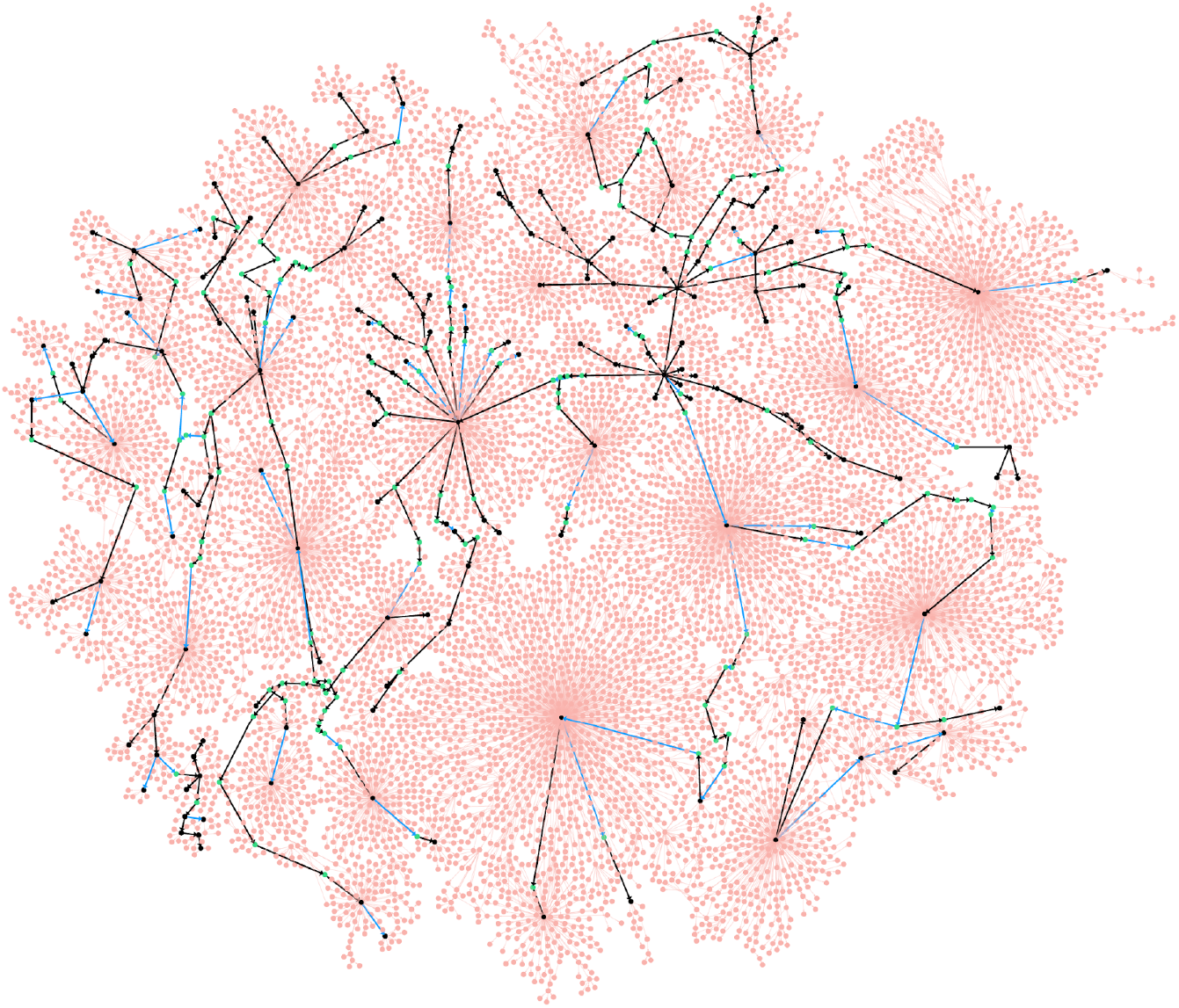
An MST of the largest Hamming graph in the of FLU1-4 dataset. IgEvolution iteratively deletes some low-abundance vertices (shown in pink) but retains low-abundance vertices shown in green (high-abundance vertices are shown in black). Black and blue edges represent the MST constructed after removing low-abundance vertices (an edge is pink if it is adjacent to a removed vertex). 265 black edges connect vertices with different amino acid sequences while 62 blue edges connect vertices with identical amino acid sequences (such vertices are glued in the clonal graph).

We emphasize a distinction between low-abundance leaves (that likely represent amplification/sequencing errors or dead-end evolutionary developments) and low-abundance internal vertices in an MST that are important for reconstructing antibody evolution. Since the existing repertoire construction tools do not distinguish between these two types of low-abundance vertices, they often remove all low-abundance vertices and thus make it difficult to reconstruct clonal trees. In contrast, IgEvolution iteratively removes low-abundance leaves in the MST but preserves low-abundance internal vertices so that all low-abundance vertices in the resulting clonal tree belong to paths connecting high-abundance vertices. Since there are usually multiple MSTs for a given HG, IgEvolution selects one of them as described in Supplemental Note “Prioritizing equally-weighted edges in the Hamming graph”. Supplemental Note “Performance of repertoire construction tools on clonally expanded datasets” reveals limitations of existing repertoire construction tools by benchmarking them on the FLU1-4 dataset.

### From clonal tree to clonal graph

Nucleotide sequences in different vertices of the clonal tree may correspond to identical amino acid sequences. “Gluing” vertices with identical amino acid sequences transforms a clonal tree into a *clonal graph* that may contain cycles. Each vertex *w* of a clonal graph corresponds to a protein sequence and two vertices *w* and *w’* are adjacent if there exist adjacent vertices *v* and *w’* in the clonal tree such that (i) amino acid sequence of *v* coincides with *w* and (ii) amino acid sequence of *v’* coincides with *w’.* In some cases, the clonal graph preserves the topology of the clonal tree (e.g., when gluing is limited to pairs of vertices that are connected by an edge in an MST (like for the graph in Figure 1). We say that a clonal graph was derived from a triplet of V, D, and J genes if nearly all sequences from this graph are aligned to these genes. See Supplemental Note “Types of clonal graphs”.

### From clonal graph to SHM graphs

IgEvolution aligns all sequences in the clonal graph to compare the amino acids occurring at the same position. Given a position *i*, we color each vertex of a clonal graph according to the amino acid at position *i* resulting in a *colored clonal graph*. We refer to an edge in a clonal graph as an *i-unicolored edge* if it connects two sequences with the same amino acid at position *i*.

Given an edge (*v, w*) in the graph, we define its *contraction* as substituting *v* and *w* by a single vertex *u* and adding edges connecting *u* with all vertices that *v* and *w* were connected with. Given a position *i* in a clonal graph, we define the *SHM graph* (referred to as *SHMGraph*(*i*)) as the result of contracting all *i*-unicolored edges in the clonal graph. We define the *multiplicity* of a vertex in the SHM graph as the number of vertices in the clonal graph that were contracted into this vertex. We classify a vertex of an SHM graph as a *low-multiplicity* vertex if its multiplicity does not exceed a threshold *multSHM* (see Supplemental Note “IgEvolution parameters”). IgEvolution removes low-multiplicity leaves from the SHM graphs and refers to the remaining edges as *salient SHM*s. In contrast to the usual approach that simply counts the number of different amino acids at each position, this approach allows one to identify positions accumulating many SHMs.

Figure 2 shows the colored clonal graphs for the largest lineage in the FLU1-4 dataset (with vertices colored according to amino acids) at position 34 (located in FR2) and 57 (located in CDR2) as well as SHM graphs *SHMGraph*(34) and *SHMGraph*(57). Both positions are characterized by several amino acids (4 amino acids at position 34 and 7 amino acids at position 57). However, as Figure 2 illustrates, SHMs affecting position 34 correspond to edges leading to low-multiplicity leaves that likely represent dead-end evolutionary development in this lineage. Thus, position 34 is characterized by a single dominant amino acid (Met) and zero salient SHMs. In contrast, position 57 is characterized by 5 dominant amino acids and 19 salient SHMs. Thus, using SHM graphs, we can classify position 34 as conservative and position 57 as *highly mutable*.

**Figure 2.**
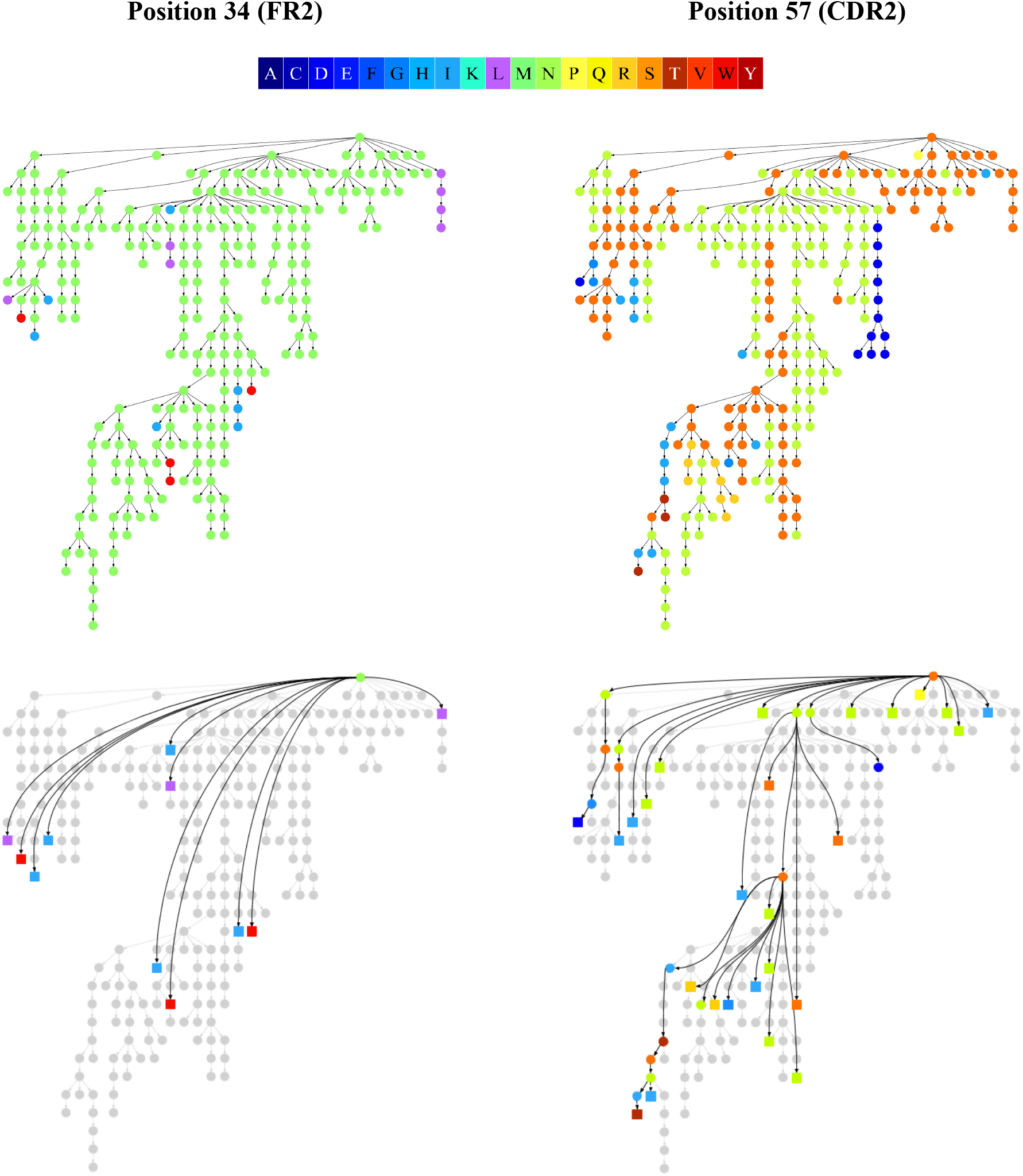
Colored clonal graphs and corresponding SHMs graphs for amino acid position 34 in FR2 (left) and amino acid position 57 in CDR2 (right) in *FluGraph*. The color scale for twenty amino acids is shown at the top. Every vertex in the SHM graph corresponds to a unicolored component in the clonal graph. High-multiplicity (low-multiplicity) vertices of the SHM graphs are shown as circles (squares).

In general, we expect that positions participating in antigen binding are characterized by higher diversity of SHMs (as they reflect multiple optimizing steps in an antibody evolution) as compared to conservative positions that are mostly responsible for maintaining antibody’s stability. We thus define the *mutability* of a position in a clonal graph as the fraction of salient SHMs at this position among all salient SHMs.

### The challenge of SHM analysis

Given an antibody repertoire, a still unsolved problem is to figure out which SHMs in this repertoire are important for antibody-antigen binding and which ones represent “dead-ends” in an evolutionary development of a repertoire. The SHMs for a single antibody are defined as all differences between its sequence and the closest germline gene. Each such SHM is defined by a pair: the nucleotide (amino acid) position in the germline gene and the nucleotide (amino acid) corresponding to this position in the antibody (we refer to this approach as *individual SHM counting*). Previous studies assumed that all antibodies in a clonal lineage have the same length and defined the *SHM-set* for a *clonal lineage* as all such distinct pairs across all antibodies in a clonal lineage (we refer to this approach as *lineage SHM counting*). Note that individual and clonal SHM counting usually do not account for SHMs in CDR3s since it is difficult to infer which D gene and non-genomic insertions contributed to a given antibody.

Many studies have analyzed the distributions of the number of SHMs across all individual antibodies (Wendel et al., 2017; Lee et al., 2019) or across all clonal lineages (Ellebedy et al., 2016; Nielsen et al., 2019; Horns et al., 2019) in an antibody repertoire. These approaches, while valuable, face the difficulty of finding the closest D genes and identifying non-genomic insertions, leading to complications in deriving SHMs in CDR3s. Moreover, they do not account for the fact that SHMs arise on the edges of the clonal trees. As a result, they deviate from the classical approach to mutation analysis accepted in modern phylogenetics, corrupt real multiplicities of SHMs, and make it difficult to analyze convergent SHMs that shed light on functionally important antibodies.

Stern et al., 2014 and Magri et al., 2017 argued that the lineage SHM counting approach is deficient and instead analyzed SHMs defined by all edges of a clonal tree. However, since it is not clear how to derive a repertoire for a clonal tree reconstruction, these studies used stringent thresholds for repertoire construction that resulted in rather small clonal lineages. Here, we describe a large-scale *clonal graph based SHM counting* across multiple clonal lineages and its applications to analyzing SHMs that affect antibody-antigen binding.

We computed the mutability of all positions in the amino acid sequences of the largest clonal graph of the FLU1-4 dataset (referred below as *FluGraph*) and compared it with the mutability computed using individual and lineage SHM counting approaches. Figure 3 shows that each approach reveals a different set of highly mutable positions. In the case of individual SHM counting, these positions correspond to early SHMs occurring in many sequences from the lineage and thus counted as SHMs in each of them. The lineage SHM counting approach provides a better estimate of mutability but fails to identify convergent SHMs and distinguish conservative positions (e.g., position 34 in Figure 2) from positions with high amino acid diversity (e.g., position 57 in Figure 2), thus overestimating the mutability of conservative sites.

**Figure 3.**
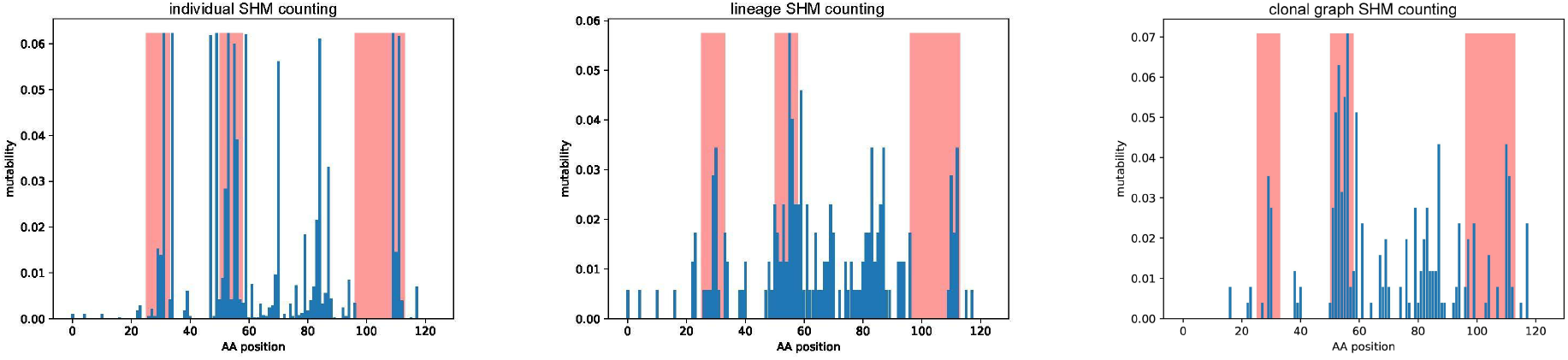
Individual counting (left), lineage counting (middle), and clonal graph counting (right) applied to *FluGraph*. A bar at position *i* shows the mutability of position *i* in *FluGraph*. Red bars correspond to positions of CDRs according to the IMGT notation.

## Results

### Datasets

We used IgEvolution to analyze the following immunosequencing datasets (Table S1):
- FLU 1: antibody repertoires of four humans taken at 7th, 14th, and 28th days after a flu vaccination (PRJNA324093 project, Ellebedy et al., 2016).
- FLU 2: antibody repertoires of four humans taken after a flu vaccination (PRJNA512111 project, Horns et al., 2019).
- INTESTINAL: intestinal antibody repertoires from four humans (PRJNA355402 project, Magri et al., 2017).

We combined all samples corresponding to the same individual (see Supplemental Note “Immunosequencing datasets”) and launched IgEvolution on each combined dataset. A lineage is classified as *large* if it includes at least *minReadNumber* reads (the default value *minReadNumber* = 50). A clonal graph is classified as *large* if it includes at least *minVertexNumber* vertices (the default value *minVertexNumber* = 50). Table S2 shows that the number of clonal lineages is typically ~10 times smaller than the number of distinct reads in a Rep-seq dataset and that large clonal lineages form only 1–8% of the total number of clonal lineages (across all datasets). The clonal tree-based error correction significantly reduces the number of sequences and simplifies the choice of interesting clonal lineages (the average number of large clonal graphs across all datasets is 7). Supplemental Note “Clonal analysis of rat antibody repertoires” describes applications of IgEvolution to ten rat repertoires analyzed in VanDuijn et al., 2017).

### Mutability of CDRs and FRs

Mutabilities of positions in CDRs (corresponding to most antigen-binding sites) across 78 large clonal graphs vary by three orders of magnitude from 0.0005 to 0.26 with the median *mut*_med_ = 0.031. We say that a position *i* in a clonal graph is *highly mutable* if its mutability exceeds *mut*_med_(see Supplemental Note “Finding highly mutable positions”).

The *mutability of a region* is defined as the average mutability of positions in this region. Figure 4 shows the distribution of mutabilities across CDRs and FRs in 78 large clonal graphs and reveals that, as expected, the average mutability of CDRs (0.015) is higher than the average mutability of FRs (0.005). Somewhat surprisingly, CDR1s and CDR2s have higher average mutabilities than CDR3s and FR3s has similar mutability to CDR3, suggesting that there exist alternative antigen-binding sites in FR3.

**Figure 4.**
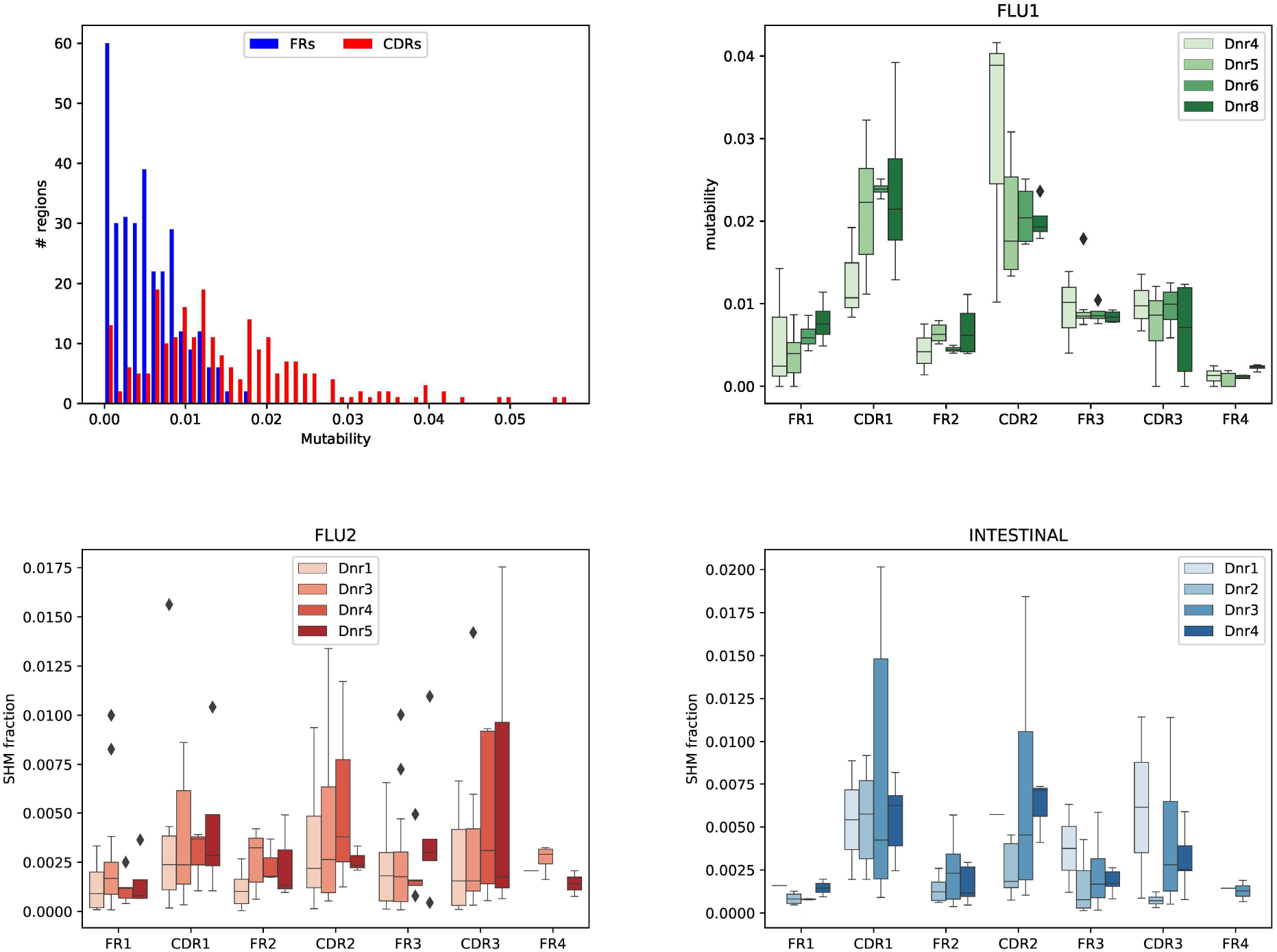
Mutability of FRs and CDRs. (Top left) Mutability of FRs and CDRs across 78 large clonal graphs. (Top right, bottom left, bottom right) Mutability of CDR1–CDR3 and FR1–FR4 regions in the FLU1, FLU2, and INTESTINAL datasets. Since the reads from the FLU1 and INTESTINAL datasets do not cover the first 50 nucleotides of FR1s, we normalized mutability by the length of the FR1 segment present in these datasets.

### Analysis of antibody-antigen 3D structures

Figure 3 suggests that some antibodies undergo extensive optimization at positions outside the conventional antigen binding sites. We thus analyzed three-dimensional structures of all 745 crystallized human antibody-antigen complexes from the SAbDab database (Dunbar et al., 2014) to identify non-conventional antigen binding sites and to check whether they correlate with highly mutable positions revealed by IgEvolution. For each complex, we identified positions in the variable region of the heavy chain contacting the antigen (*binding sites*) and the light chain (*contact sites*) as described in Supplemental Note “Finding binding and contact sites in antibody-antigen complexes” (Figure 5).

**Figure 5.**
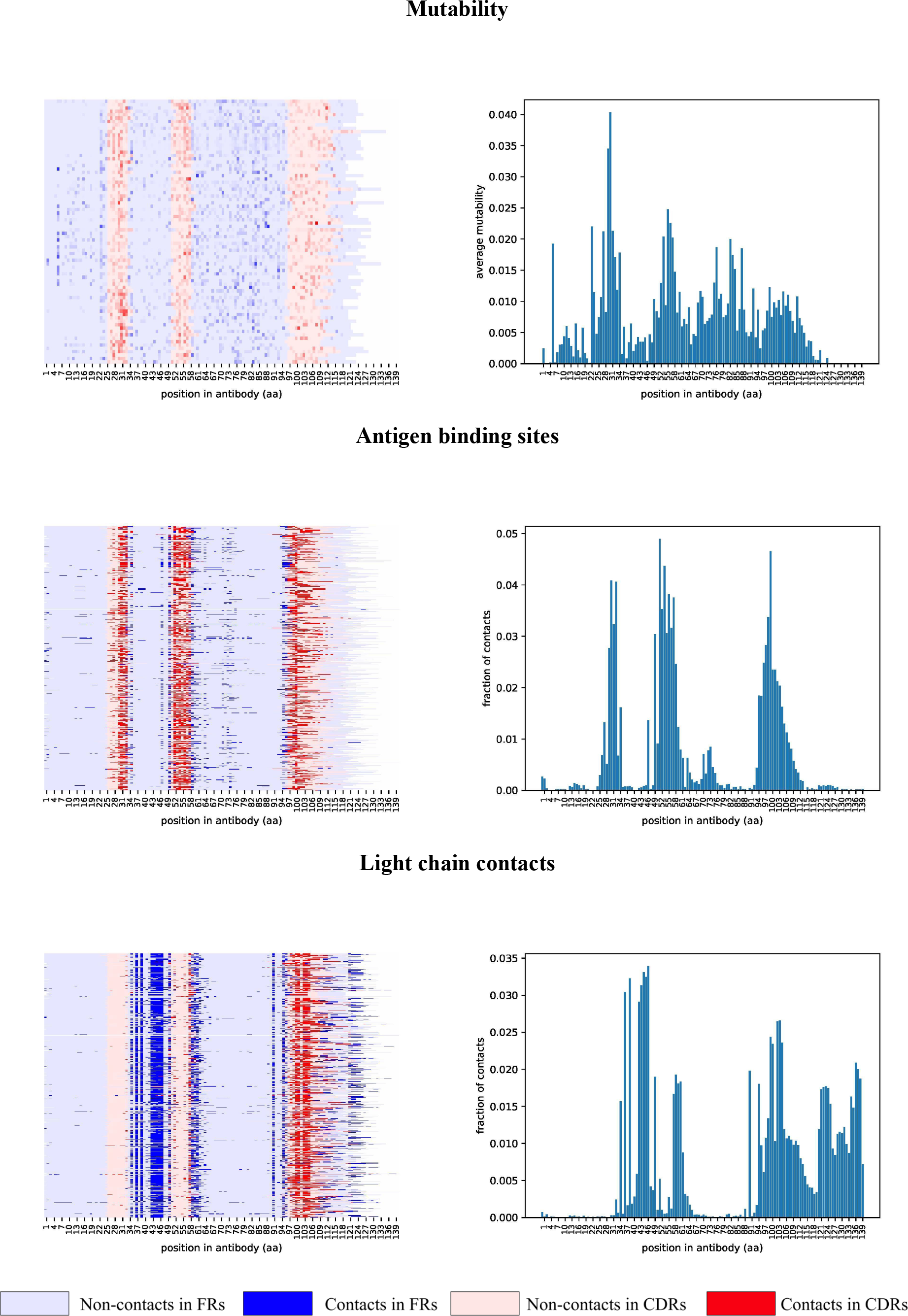
Maps of mutability (top), antigen contacts (middle), and light chain contacts (bottom). Mutability map is computed using 78 large clonal graphs from all four analyzed immunosequencing datasets. Maps of contacts were computed using 745 heavy chain antibody sequences extracted from antibody-antigen complexes. Sequences of heavy chains are aligned by the first position and cropped after position 140. Each heavy chain antibody sequence corresponds to a row in the map. Positions of FR and CDR regions are derived according to the IMGT notation and are shown in blue and red, respectively. (Top) Mutability of a position is shown according to a color palette. (Middle, bottom) Positions corresponding to contacts are shown in a darker color.

As Figure 5 illustrates, binding and contact sites are often conserved among multiple antibodies. In contrast to CDR1 and CDR2 that contain binding sites (and hardly any contact sites), CDR3 contains both binding and contact sites. In contrast to FR1 that has hardly any contact sites, FR2 has many contact sites. Also the first 1–3 and the last 1–2 positions of FR2 often represent binding sites, suggesting that the IMGT definition of CDR1 and CDR2 misses some important positions. The first 1–8 and the last 1–7 positions of FR3 also correspond to both binding sites (extending CDR2 and CDR3) and contact sites. Also, many FR3s contain non-conventional binding sites centered at distance 16 from the beginning of FR3 (this region named *CDR4* in Kirik et al., 2017). The last position of FR4 corresponds to contact sites that are extended in constant regions.

Since the lengths of CDRs and FRs vary across different antibodies, below we explain how to select *representative lengths* for each CDR/FR and analyze mutability of positions in each CDR/FR region along its length.

### Highly mutable positions in clonal graphs correlate with binding and contact sites

To check whether highly mutable positions in clonal graphs correlate with binding and contact sites in antibodies, we limit the analysis to regions (FR1, CDR1, FR2, CDR2, FR3, and FR4) of the same *typical* length. For example, the FR1s has a typical length 25 aa in 74% (78%) of the crystallized antibodies (the clonal graphs). 86%, 99%, 77%, 96%, and 95% of CDR1, FR2s, CDR2, FR3s, and FR4 have lengths 8, 17, 8, 38, and 11 aa across all heavy chain sequences in the crystallized antibodies. Below we refer to such equally-sized regions as *typical regions*. Note that the concept of typical length is not applicable to CDR3.

Below we describe the joint analysis of all 3D antibody structures and clonal graphs for typical regions (FR1, CDR1, FR2, CDR2, FR3, and FR4). For each position in each typical region, we computed (i) the percentage of the structures where this position corresponds to a binding or/and a contact site, and (ii) the percentage of clonal graphs where this position is highly mutable. On average, a position is highly mutable in *percent*_*med*_ = 8% of clonal graphs.

Finally, we analyze salient SHMs from all positions in each typical region. Each such SHM is defined by a pair: its position within a typical region and a new amino acid at this position. A *multiplicity* of a SHM is the number of times its pair appears in clonal graphs. We define a SHM as *convergent* if its multiplicity exceeds *mult*_*min*_ (the default *mult*_*min*_= 12 is the upper quartile of all SHM multiplicities).

### Analyzing highly mutable positions in CDR1

Figures 5 and 6 illustrate that some positions within typical CDR1 (8 aa long) have a large contribution to antigen binding (e.g., position 6 is a binding site in 53% of antibodies) while some positions have a small contribution (e.g., each of positions 1, 2, and 4 is a binding site in less than 8% of antibodies). Figure 6 also shows that non-binding positions 1, 2, and 4 in CDR1 rarely mutate in the clonal graphs (these positions are classified as highly mutable in only 4 – 8% of clonal graphs), while the most frequent binding position 6 is also the most mutable among all positions (represents a highly mutable position in 36% of clonal graph). The binding sites with the highest mutability in CDR1 (positions 3, 5, and 6) feature the largest number of distinct SHMs (varying from 43 to 64).

**Figure 6.**
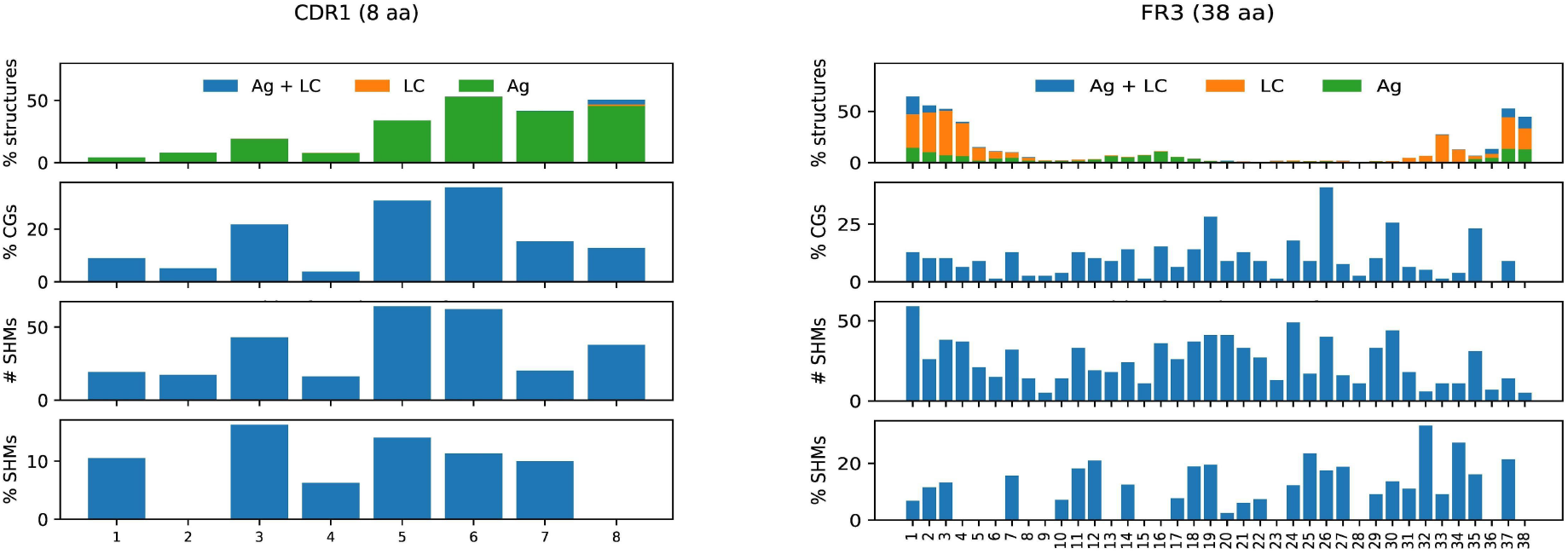
Characteristics of typical CDR1 (left) and FR3 (right). For each region, we computed four plots described below. (i) A bar at a position within a typical region shows the percentage of binding sites (green), contact sites (orange), and both binding and contact sites (blue) at this position. (ii) A bar at a position shows the percentage of clonal graphs where this position is highly mutable. (iii) A bar at a position shows the number of distinct SHMs appearing at this position across all large clonal graphs. (iv) A bar at a position shows the percentage of convergent SHMs appearing at this position across all large clonal graphs. The percentage is computed with respect to the number of distinct SHMs appearing at this position.

Figures 6 and S2 shows the results of the same analysis applied to other regions except for CDR3 (since there is no typical length for CDR3s) and FR4 (since no clonal graph has highly mutable positions in FR4). This analysis revealed interesting properties that shed light on functions of various regions of antibodies (Supplemental Note “Analyzing highly mutable positions in typical FR1, FR2, and CDR2”). In particular, the first and the last positions of FR2 and FR3 have high mutability in clonal graphs thus likely extending the bounds of CDR1 and CDR2.

### Analyzing highly mutable positions in FR3

We partitioned a typical FR3 region into four subregions: (i) positions 1–8 with frequent contact sites in 3–39% of 3D structures; (ii) positions 9–18 with non-conventional binding sites in 2–11% of 3D structures; (iii) positions 19–31 with hardly any contact sites in 3D structures; and (iv) positions 32–38 with frequent contact sites in 3–31% of 3D structures. Interestingly, the non-contacting subregion (iii) has high mutability in the largest percent of clonal graphs among all subregions. Only 9 out of 13 positions have high mutability in more than *perc*_*med*_ (9–41%) of clonal graphs. These positions also contain many convergent SHMs. While the role of these positions is unclear, we assume that the abundance of the convergent SHMs at these positions suggests that they may contribute to the structural properties of antibodies, e.g., conformation of CDRs (Ovchinnikov et al., 2018).

### Usage of V genes in clonal lineages

The *usage* of a germline gene is defined as the fraction of reads in a repertoire aligned to this gene. Although this concept (that we refer to as the *simple usage*) proved to be useful (Wendel et al., 2017; Lee et al., 2019), it does not accurately estimate the number of VDJ recombinations that include a given germline gene, particularly for stimulated repertoires with expanded clonal lineages. Since each clonal lineage reveals a single VDJ recombination, we define the *clonal usage* of a germline gene as the fraction of clonal graphs derived from this gene (Figure 7). Filtering trivial clonal graphs allows us to lower the impact of V genes coming from background B cells and detect V genes that are important for antigen specific immune response (Figure 7). We found that three most used V genes in the FLU1 dataset are among the five most used V genes for the FLU2 dataset: IGHV3-7 (16% and 9% of clonal graphs in FLU1 and FLU2, respectively), IGHV4-39 (13% and 6%), and IGHV1-69 (7% and 15%). In contrast, the INTESTINAL datasets have completely different most used V genes (IGHV5-51 (26%) and IGHV4-59 (17%)), suggesting a correlation between most used V genes and antigens.

**Figure 7.**
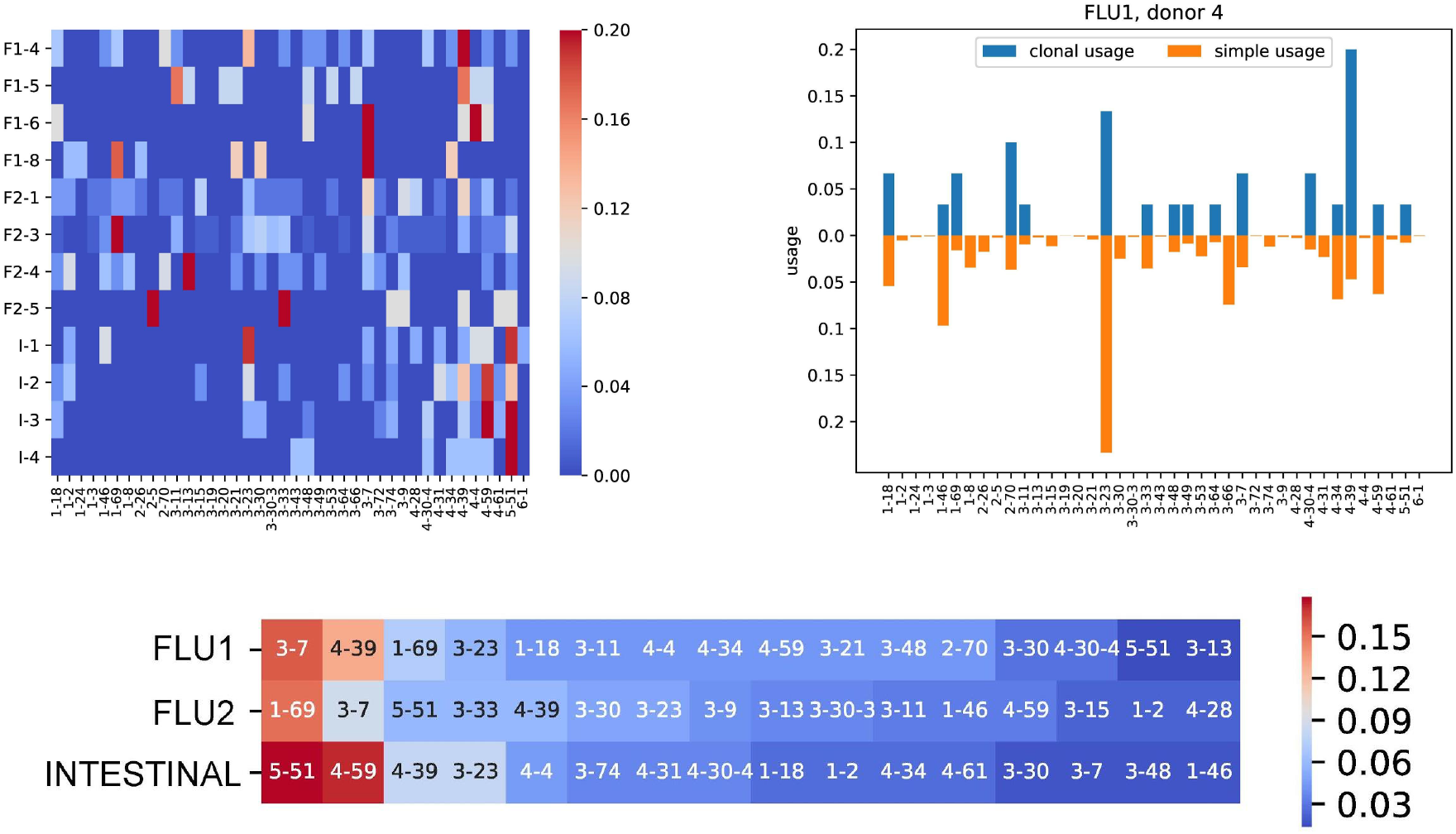
Clonal usage of V genes. Clonal usage is computed based on clonal graphs with at least 2 vertices. (Top left) Clonal usage of V genes across FLU1 (F1), FLU2 (F2), and INTESTINAL (I) datasets. (Top right) Clonal (blue) and simple (orange) usages of V genes for the FLU1-4 dataset. Usage values are shown only for V genes from the left plot. (Bottom) Heatmap of V genes in FLU1, FLU2, and INTESTINAL dataset ordered in the decreasing order of their clonal usage. The color of a cell represents the fraction of clonal graphs derived from corresponding V gene.

Clonal usage also reveals the differences between repertoires developed in different individuals in response to the same antigen. Figure 7 shows the clonal usage for the FLU1 dataset (flu vaccination, hemagglutinin specific B cells) and illustrates that IGHV1-69 (that was implicated in the immune response to hemagglutinin in previous studies) is heavily used by donors 4 and 8 (clonal usage 7% and 18%, respectively), but is not used by donors 5 and 6 (clonal usage is 0% in both donors 5 and 6).

### Different alleles of IGHV1-69 shape flu-specific responses

The FLU1 datasets consist of receptor sequences specific to hemagglutinin (HA), one of the antigens of the flu virus. Recent studies showed that many HA-neutralizing antibodies use IGHV1-69 and that genomics variations in IGHV1-69 influence the binding ability of corresponding antibodies (Lingwood et al., 2012; Avnir et al., 2016). The IMGT database includes 14 alleles of the IGHV1-69 gene (Figure 8), including *F-alleles* (with Phe at the amino acid position 55) and *L-alleles* (with Leu at the same position). Lingwood et al., 2012 demonstrated that the F-alleles result in antibodies with higher affinity to HA as compared to antibodies derived from the L-alleles. Avnir et al., 2016 showed that the usage of L-alleles in HA-stimulated antibody repertoires is much lower compared to F-alleles.

**Figure 8.**
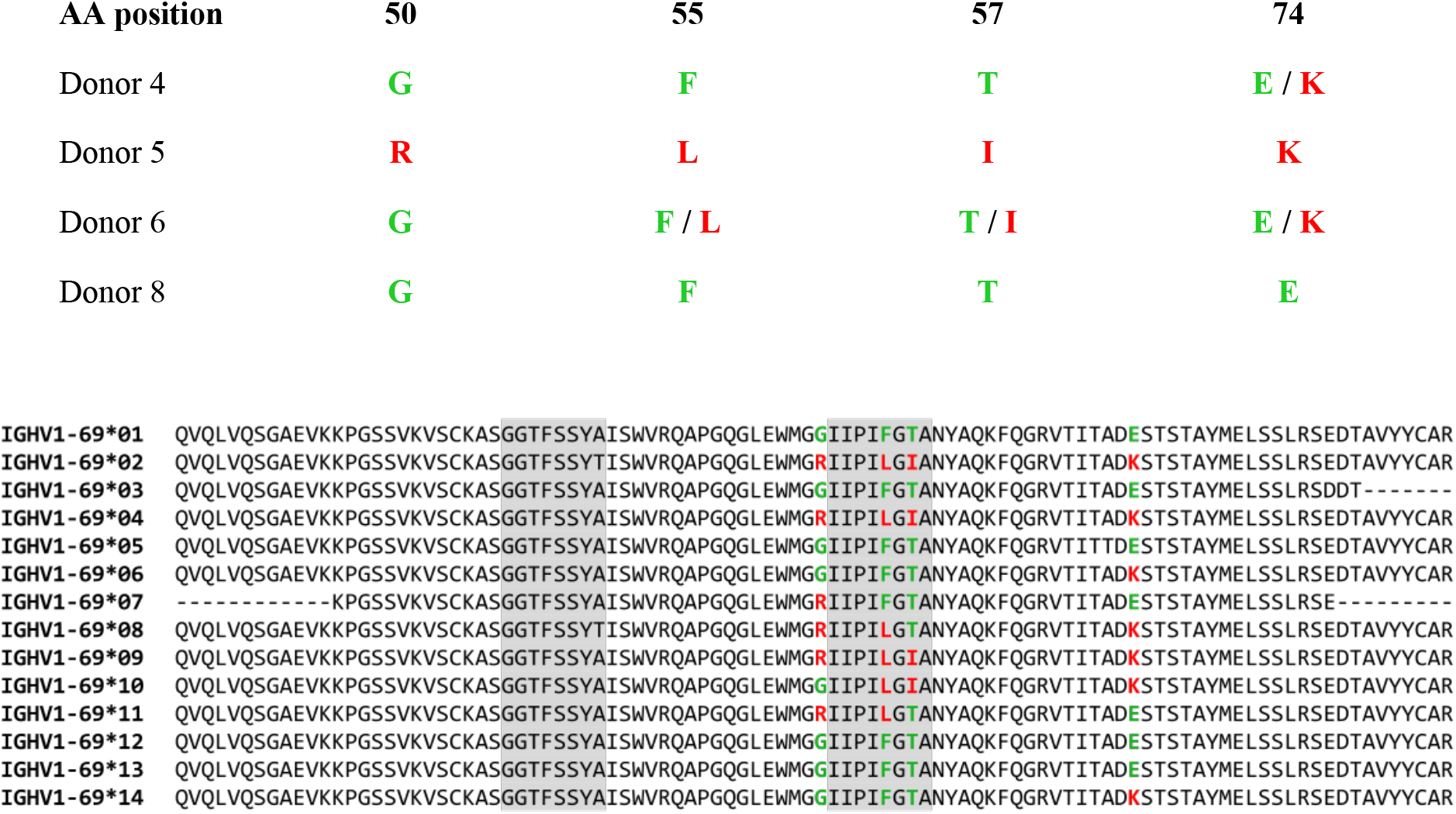
Germline variations of the IGHV1-69 gene in four individuals in the FLU1 dataset. (Top) A table shows amino acids at four positions selected as candidates for genomic variations. (Bottom) Amino acid sequences of 14 known alleles of IGHV1-69. Dominant and recessive amino acids corresponding to the detected germline differences are shown in green and red, respectively. Grey substrings highlight CDR1 and CDR2.

### Analysis of IGHV1-69 alleles in the FLU1 dataset

To derive the IGHV1-69 alleles of FLU1 donors, we analyzed naïve datasets corresponding to these donors (Supplemental Note “Test immunosequencing datasets”). For each naïve dataset, we collapsed reads with identical CDR3s, aligned them against known human V genes (we used the first alleles only), and identified reads aligned to IGHV1-69. Afterwards, we selected candidates for germline variations in IGHV1-69 as positions in the alignment where at least 40% of aligned reads vote for a non-germline nucleotide. As a result, we selected four nucleotide positions: 148, 163, 170, and 220 corresponding to amino acid at positions 50, 55, 57, and 74, respectively. Afterwards, we analyzed two alternative amino acids at four selected positions. Figure 8 shows that the found positions are confirmed by known alleles of IGHV1-69.

Figure 8 shows that donors 4 and 8 have F-alleles (both donors have Phe at amino acid position 55), donor 5 has L-allele, and donor 6 is heterozygous with both F- and L-alleles. Amino acids at three other detected positions also separate donors 4 and 8 from donors 5 and 6. We refer to amino acids at positions 50, 55, 57, and 74 inferred from the naive datasets for both donors 4 and 8 as *dominant* and to alternative amino acids at these positions as *recessive*. This analysis revealed that amino acids G (R), F (L), T (I), and E (K) at positions 50, 55, 57, and 74 are dominant (recessive).

Since donors 4 and 8 have F-alleles (that provide effective flu response), donor 5 has L-alleles (that lack effective flu response), and donor 6 has both F- and L-alleles, we expect high clonal usage of IGHV1-69 in donors 4 and 8 and low clonal usage in donors 5 and 6 (Lingwood et al., 2012, Avnir et al., 2016). Indeed, IGHV1-69 is utilized in clonal graphs of donors 4 and 8 but missing in clonal graphs of donors 5 and 6 (Figure 7). Interestingly, our analysis reveals that, in addition to position 55 in IGHV1-69 (that has known impact on contribution to response against HA), positions 50, 57, and 74 (that also separate donors 4 and 8 from donors 5 and 6) may also represent candidate positions affecting the immune response against HA.

### Clonal graphs reveal the role of IGHV1-69 alleles

Clonal graphs derived from IGHV1-69 may shed light on the role of germline variations in immune response. If an amino acid is important for antigen binding or maintaining the antibody structure, we expect to see a strong selection against changing this amino acid during immune response. If an amino acid is not important, then it will likely be substituted by another amino acid during clonal development. We thus conjecture that some dominant amino acids of IGHV1-69 do not mutated in clonal graphs derived from this gene in the FLU datasets. Finding such amino acids helps us to analyze associations between alleles of V genes and specific immune response.

To check this hypothesis, we selected clonal graphs derived from IGHV1-69 for FLU1-4 and FLU1-8 datasets that represent individuals with dominant alleles. For each selected clonal graph, we analyzed positions 50 (Gly / Arg), 55 (Phe / Leu), 57 (Thr / Ile), and 74 (Lys / Glu) (Figure 8). Figure S3 shows that Gly at position 50 and Thr at position 57 are preserved in all selected clonal graphs. Phe at position 55 is preserved in all clonal graphs except for one where it is partially substituted with Ser. Lys at position 74 is preserved in all clonal graphs except for one where it is substituted with Thr.

To extend this analysis, we analyzed all 33 clonal graphs derived from the IGHV1-69 in the FLU2 datasets. Although the FLU2 datasets were not sorted against HA, they were sequenced after flu vaccination and thus are likely to include some HA-stimulated lineages. Figure S4 shows amino acid content in the selected clonal graphs at positions 50, 55, 57, and 74 and confirms that amino acids Gly, Phe, and Thr are present in most clonal graphs at positions 50, 55, and 57, respectively. Thus, we conjecture that positions 50, 55, and 57 in IGHV1-69 are associated with the immune response to flu.

However, for position 55, we found that the largest clonal graph in the FLU2 datasets has similar fractions of Phe and Leu (Figure 9). This case suggests that sequences containing Phe and Leu evolve independently and thus the absence of Phe does not necessarily lead to a complete loss of binding properties of the antibody.

**Figure 9.**
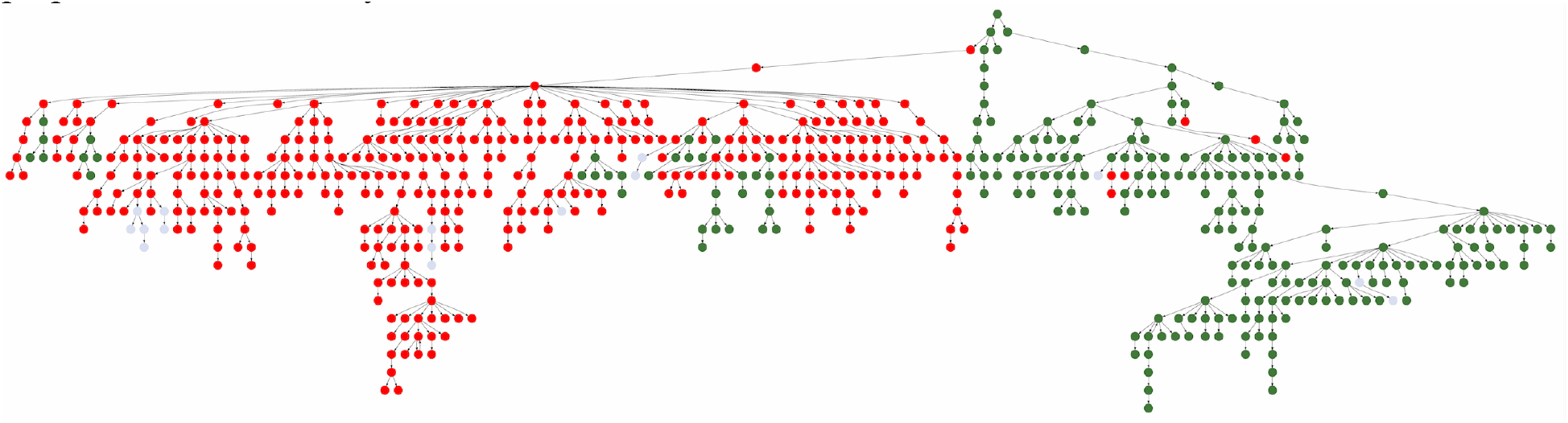
The largest clonal graph derived from IGHV1-69 in the FLU2-3 dataset. Vertices of the graph are colored according to amino acids presented at the position 55 (F – green, L – red, other amino acids – grey).

## Discussion

Although clonal analysis of immunosequencing data has important applications in monitoring treatment, measuring vaccine efficacy, and designing antibody drugs, the existing tools for constructing clonal trees have limitations. Moreover, it remains unclear how to pre-process immunosequencing datasets for follow-up clonal tree reconstruction: constructing clonal trees on all reads faces the challenge of dealing with high error rates in Rep-seq dataset (Lee et al., 2017), while constructing clonal trees on error-corrected antibody repertoires faces the challenge of constructing accurate antibody repertoires in the case of stimulated Rep-seq datasets (Shlemov et al., 2017). Although the latter challenge can potentially be addressed by molecular barcoding techniques (Horns et al., 2017), these techniques are still rarely used (most publicly available Rep-seq datasets do not contain molecular barcodes). As a result, clonal analysis of biomedically important immunosequencing samples (such as antibody repertoires after flu vaccination) remains challenging. IgEvolution addresses this challenge by developing an algorithm for simultaneous error correction and clonal reconstruction of antibody repertoires. Extending previous approaches for clonal reconstruction (Horns et al., 2017; Lee et al., 2017), IgEvolution decomposes raw Rep-seq reads into clonal lineages, constructs the minimum spanning tree for each clonal lineage, and iteratively identifies erroneous sequences located in leaves. We applied IgEvolution to analyze multiple immunosequencing datasets, described an approach to identify salient SHMs based on clonal tree analysis, and demonstrated that it reveals novel interesting features of antigen-specific antibody repertoires.

IgEvolution identified many large clonal graphs that we further analyzed to reveal various characteristics of structural regions (CDRs and FRs) in antibodies. To understand the properties of SHMs in these regions, we analyzed all crystallized human antibody-antigen complexes in the SAbDab database (Dunbar et al., 2014) and identified positions of the contacts of the heavy chain with the light chain (LC) and the antigen (Ag). This analysis revealed that structural regions of VDJ sequence have conservative functions: FR1 rarely contacts LC and Ag; CDR1 and CDR2 contact Ag; FR2 contacts LC; CDR3 contacts both LC and Ag; and FR4 contacts LC. For FR3, we revealed four subregions with different properties: (i) LC contacts; (ii) non-conventional Ag binding site (present in ~10% of antibodies); (iii) non-contacting subregion; and (iv) LC contacts.

We compared the binding and contact positions (based on analysis of crystallized 3D structures) with highly mutable positions derived from the clonal graphs and found that these two sets of positions are well correlated in CDR1 and CDR2. We also demonstrated that regions responsible for light chain binding (FR2, FR4, and prefix/suffix of FR3) rarely mutate with notable exception of the highly mutable first/last positions of FR2 and FR3 that correspond to binding sites and extend bounds of CDR1 and CDR2 as defined by traditional similarity-based tools (Kabat et al., 1979; Chothia and Lesk, 1987; Giudicelli et al., 1997). Our analysis of rat antibodies led to similar results with respect to highly mutable positions in various CDRs and FRs.

Since Rep-seq dataset that include antibodies with known 3D structures are currently unavailable, our analysis of clonal graphs and crystallized Ab-Ag complexes was performed on unrelated datasets. However, even this imperfect analysis revealed that it would be beneficial to resolve three-dimensional structure of antibody-antigen complexes and trace their evolutionary development using Rep-seq data. We hope that such paired datasets will emerge in the future and shed light on the role of SHMs in CDRs and FRs (especially FR3). At the same time, we believe that the results described in this paper can already be used in the design of antibody drugs and humanization of non-human immunoglobulins.

We also analyzed the usage of V genes in stimulated Rep-seq datasets and showed that it correlates with genomic variations of V genes. We analyzed the usage of various alleles of IGHV1-69 (V gene that is associated with hemagglutinin (HA) specific antibodies) and demonstrated that F-alleles of IGHV1-69 are used in HA-specific clonal graphs, thus providing independent support for a recent discovery of correlations between specific alleles and vaccine responses (Avnir et al., 2016). We also showed that amino acids that distinguish F-alleles from L-alleles of IGHV1-69 rarely mutate in HA-specific clonal graphs and thus undergo strong selection during the clonal development. We also analyzed alleles of IGHV3-11 and IGHV4-39 that are used differently in clonal graphs and found positions corresponding to genomic variations that undergo selection pressure (Supplemental Note “Alleles of IGHV3-11 and IGHV4-39 shape immunoglobulin response”). We thus suggest that clonal analysis of various alleles has a potential for identifying important genomic variations that contribute to responses to vaccination and disease. Our analysis indicates that high-throughput clonal analysis of Rep-seq datasets from antigen-exposed antibody repertoires will result in a database of putative associations between germline alleles and specific immune responses. Such database may enable personalized immunogenomics approaches by predicting efficiency of individual antibody response to a specific antigen.

## Availability of data and materials

IgEvolution is available at https://immunotools.github.io/immunotools. Results of IgEvolution on test immunosequencing datasets are available at https://immunotools.github.io/ig_evolution_results/.

## Author contributions

Y.S. implemented the IgEvolution algorithm, designed the computational experiments, and performed benchmarking. Y.S. and P.A.P. conceived the study and wrote the manuscript.

## Acknowledgements

Authors are grateful to Anton Bankevich, Siavash Mirarab, and Corey Watson for insightful comments and fruitful discussion.

