## Supplemental Materials for "IgEvolution: clonal analysis of antibody repertoires"

### Supplemental Figures

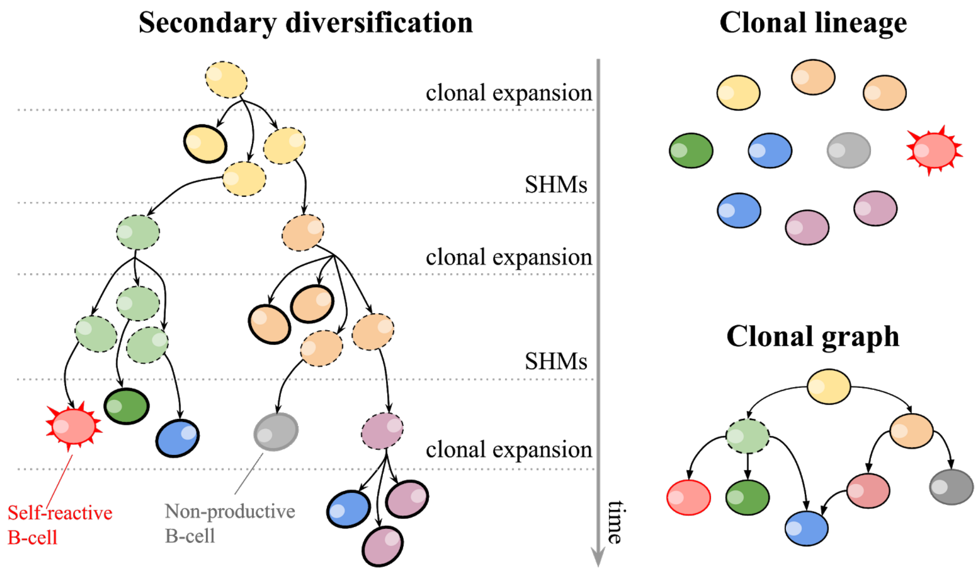


**Figure S1. Development of the antibody response as an evolutionary process.** Secondary diversification turns a single B-cell into a lineage of clonally related cells (represented as a clonal tree) through multiple rounds of clonal expansion and SHMs. SHMs may generate self-reactive B cells (shown in red) and non-productive B cells (shown in grey). Some B cells (shown as dashed vertices) might disappear during the secondary diversification. B cells that survived after the secondary diversification of a single B-cell (shown as bold vertices) form a clonal lineage. The evolutionary development of a clonal lineage can be represented as a *clonal graph*.

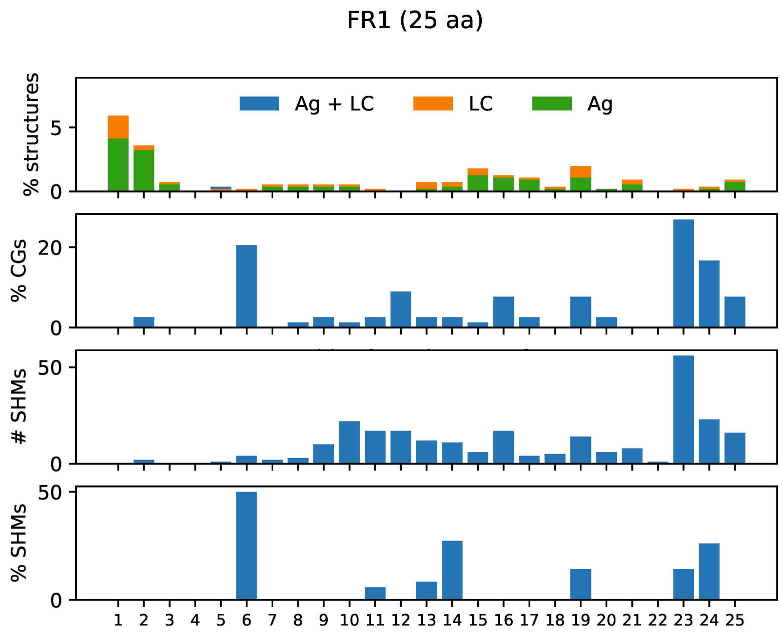

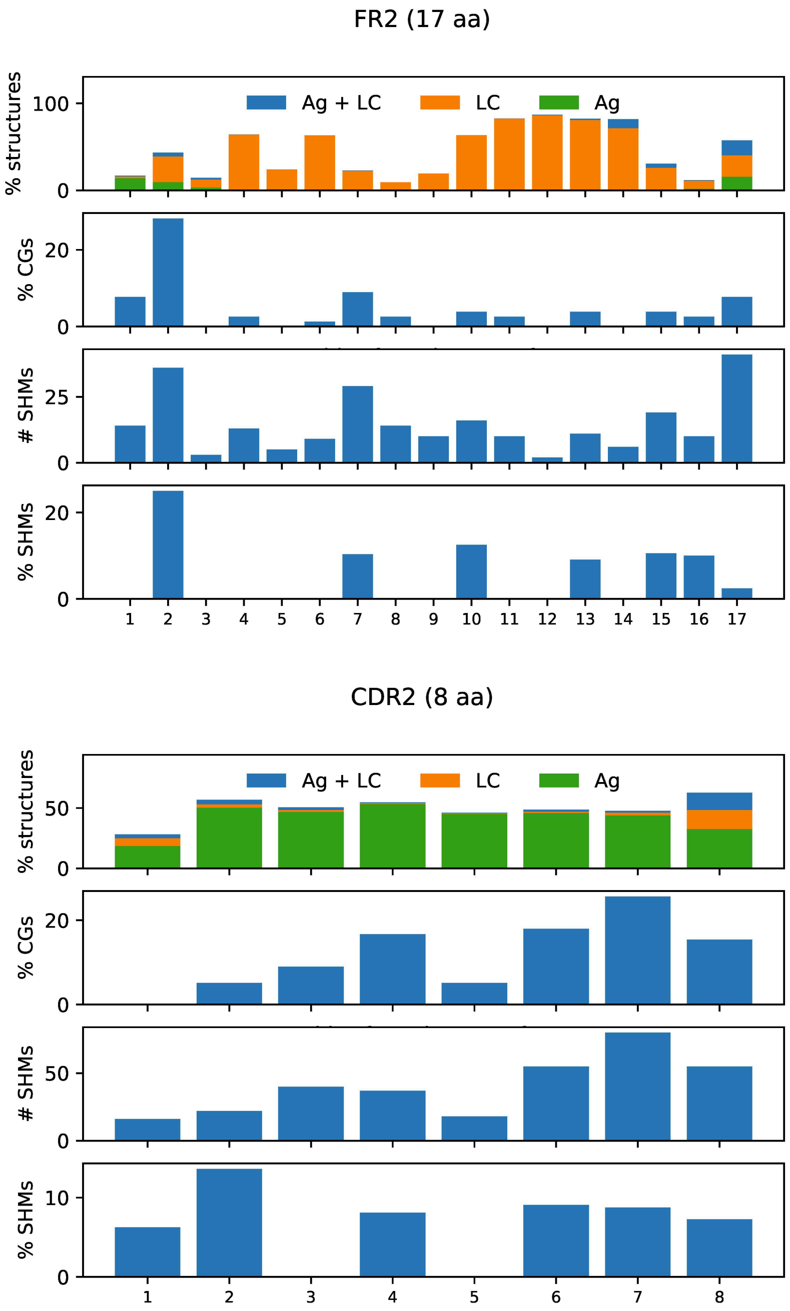


**Figure S2. Characteristics of typical FR1 (upper), FR2 (middle) and CDR2 (lower).** Description of the plots is provided in the caption of Figure 6.

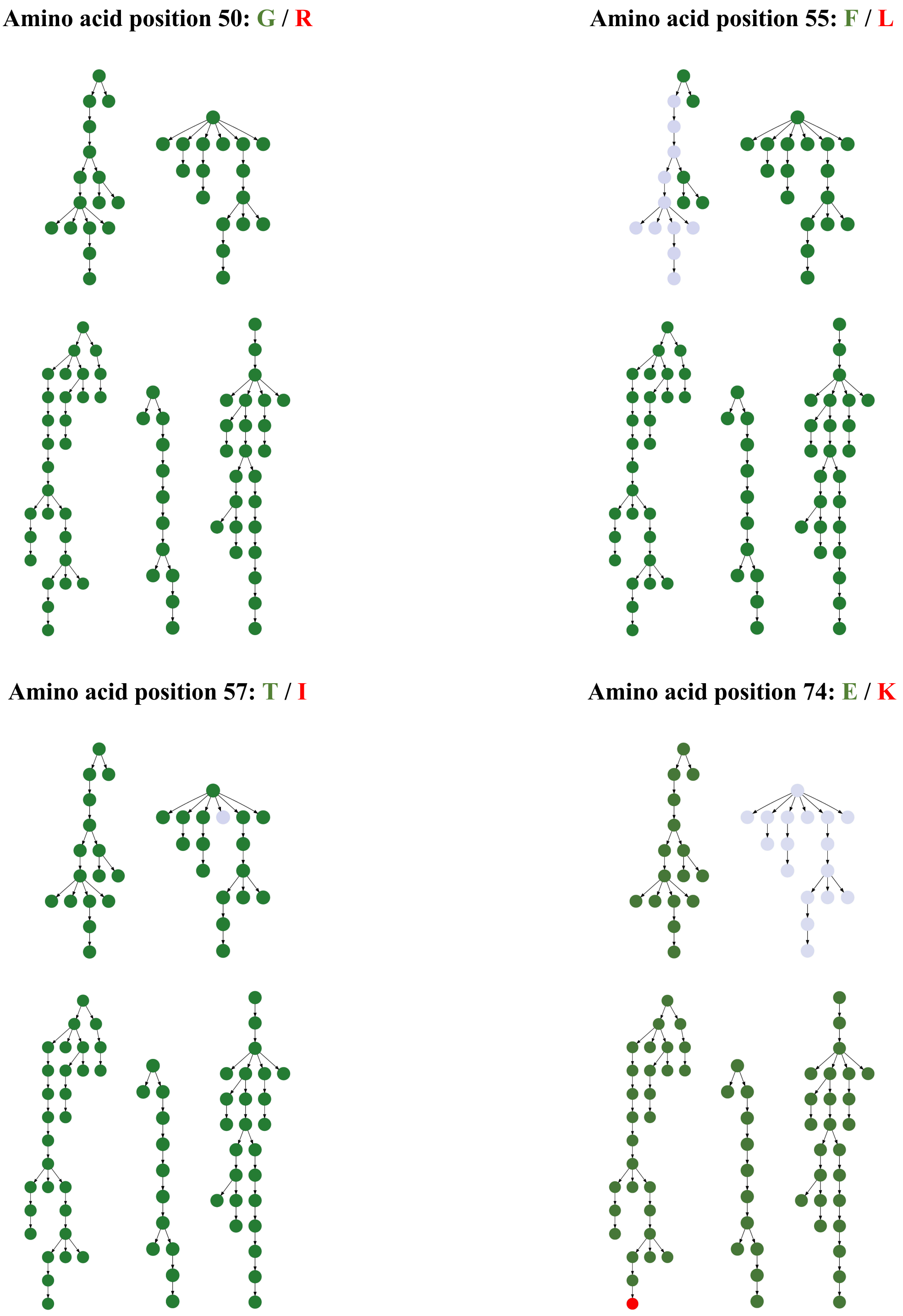


**Figure S3. Colored clonal graphs derived from IGHV1-69 in FLU1-4 and FLU1-8 datasets.** The clonal graphs are colored based on amino acids at positions 50, 55, 57, and 74. Sequences containing dominant and recessive amino acids at the corresponding position are colored in green and red, respectively. Sequences containing other amino acids are colored in grey.

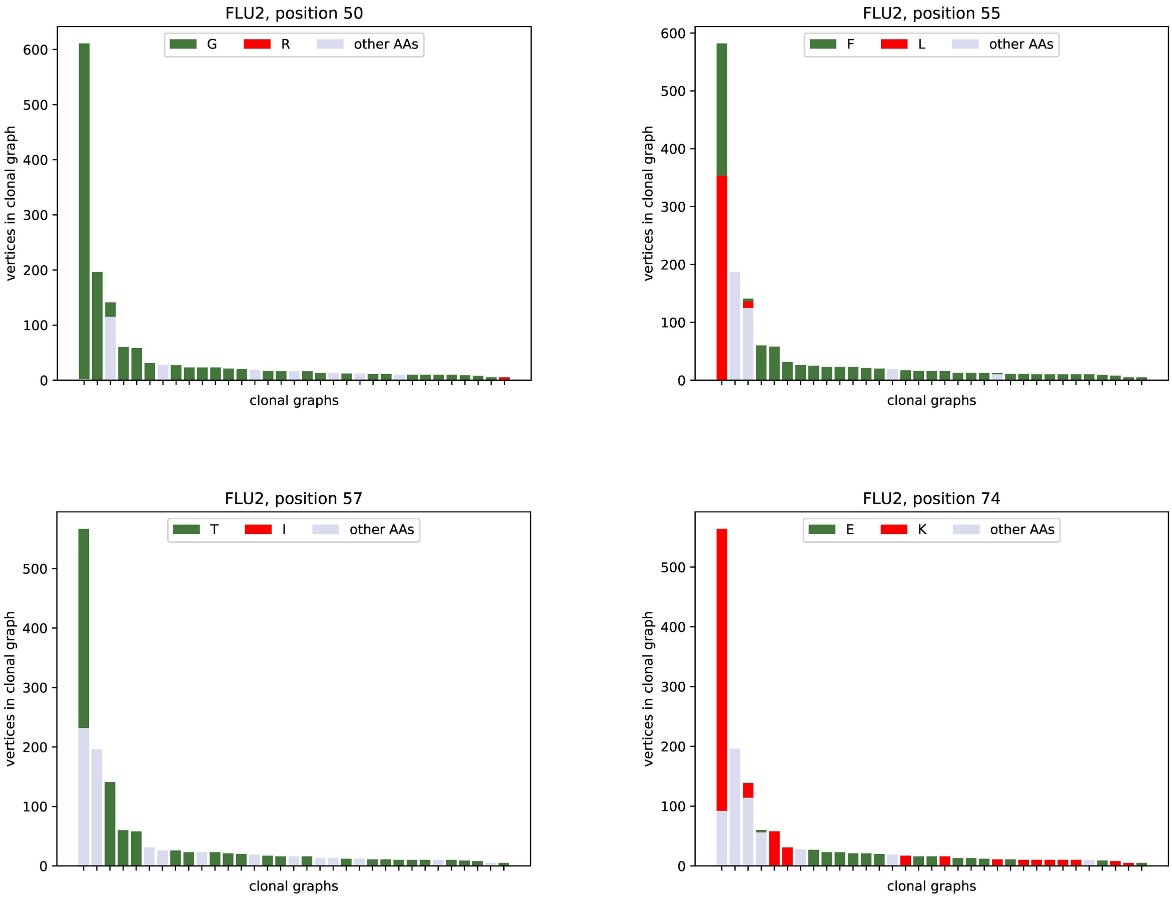


**Figure S4. Amino acid content corresponding to positions 50, 55, 57, and 74 in 33 clonal graphs derived from IGHV1-69 in the FLU2 datasets.**

### Supplemental Tables

| Group | NCBI project | Description | Cell / tissue type | # individuals |
|---|---|---|---|---|
| FLU 1 | PRJNA324093 | antibody repertoires after flu vaccination | plasma, activated, and memory HA-positive cells | 4 |
| FLU 2 | PRJNA512111 | antibody repertoires after flu vaccination | PBMC | 4 |
| INTESTINAL | PRJNA355402 | intestinal antibody repertoires | plasma and memory cells | 4 |

**Table S1. Information about human immunosequencing datasets.** HA-positive refers to cells specific to hemagglutinin (one of flu antigens).

| Dataset | Individual donor ID | # distinct reads | # clonal lineages | # (%) large clonal lineages | # large clonal graphs | # vertices /# edges in the largest clonal graph |
|---|---|---|---|---|---|---|
| FLU1 | 4 | 125,050 | 2406 | 179 (7%) | 3 | 266 / 327 |
| | 5 | 72,416 | 1119 | 73 (7%) | 7 | 425 / 599 |
| | 6 | 124,223 | 1336 | 47 (4%) | 4 | 2127 / 2746 |
| | 8 | 226,164 | 2021 | 169 (8%) | 4 | 1828 / 3206 |
| FLU2 | 1 | 1,997,159 | 118,342 | 2504 (2%) | 11 | 1790 / 2547 |
| | 3 | 1,685,626 | 105,467 | 2696 (3%) | 16 | 613 / 719 |
| | 4 | 524,408 | 65,144 | 840 (1%) | 5 | 251 / 336 |
| | 5 | 1,641,508 | 77,160 | 2839 (4%) | 5 | 147 / 199 |
| INTESTINAL | 1 | 609,292 | 40,390 | 2205 (5%) | 2 | 131 / 151 |
| | 2 | 707,302 | 50,998 | 2024 (4%) | 7 | 220 / 306 |
| | 3 | 1,131,961 | 142,674 | 2947 (2%) | 9 | 260 / 335 |
| | 4 | 472,659 | 49,862 | 1483 (3%) | 5 | 82 / 109 |

**Table S2. IgEvolution results on human immunosequencing datasets.** The “Individual donor ID” column presents the IDs used in the original papers.

### Supplemental Notes

**Constructing Hamming graphs for follow-up clonal tree reconstruction**

**Clonal lineage assignment**
**Prioritizing equally-weighted edges in the Hamming graph**

**IgEvolution limitations**

**IgEvolution parameters**

**Setting a threshold for detecting low-abundance receptor sequences**
**Finding low-multiplicity leaves in SHM graphs**
**Finding highly mutable positions**

**Performance of repertoire construction tools on clonally expanded datasets**

**Types of clonal graphs**

**Test immunosequencing datasets**

**Clonal analysis of rat antibody repertoires**

**Finding binding and contact sites in antibody-antigen complexes**

**Analyzing highly mutable positions in typical FR1, FR2, and CDR2**

**Analyzing highly mutable positions in FR1**
**Analyzing highly mutable positions in FR2**
**Analyzing highly mutable positions in CDR2**

**Alleles of IGHV3-11 and IGHV4-39 shape immunoglobulin response**

**Clonal analysis of IGHV3-11**
**Clonal analysis of IGHV4-39**

##### Supplemental Note: Constructing Hamming graphs for follow-up clonal tree reconstruction

**Clonal lineage assignment.** Since repertoire reconstruction tools attempt to identify reads derived from *identical* immunoglobulins, our HG construction algorithm (Shlemov et al., 2017) uses a rather stringent default distance threshold for defining edges in the HG (vertices are connected by an edge if the Hamming distance between them does not exceed $\tau_{IGREC} = 4$) Since clonal reconstruction tools attempt to identify reads derived from *clonally related* (rather than identical) immunoglobulins, IgEvolution increases this threshold to meet the following two conditions: (i) mutated but clonally related immunoglobulins belong to the same connected component of the HG, and (ii) clonally unrelated immunoglobulin belong to different connected components.

Since simply increasing the threshold $\tau_{IGREC}$ may join clonally unrelated immunoglobulins (and thus violate the condition (ii)), we first decompose the Rep-seq sequences into clonal lineages. IgEvolution computes a Hamming graph on CDR3s and reports connected components of the graphs as clonal lineages. To avoid connections between non-related sequences with similar short CDR3s, IgEvolution uses different thresholds for different CDR3 lengths (at most 3 mismatches for 0–30 nt long CDR3s; at most 6 mismatches for 30–60 nt long CDR3s, etc). This approach was adopted from Horns et al., 2016. Alternatively, clonal lineages can be derived using existing clonal lineage assignment tools Change-O (Gupta et al., 2014), Clonify (Briney et al., 2016), and partis (Ralph and Matsen, 2016). Afterwards, IgEvolution constructs the HG for each clonal lineage using the large default value $\tau_{IGREC} = 15$ and use the largest connected component of this graph for the MST construction. Note that the HG construction, based on substitutions only, does not account for insertions and deletions. However, the Supplemental Note "IgEvolution limitations" demonstrates that indels have only a small impact on analyzing antibody repertoires.

**Prioritizing equally-weighted edges in the Hamming graph.** A graph with unique edges-weights have a single MST but graphs with non-unique edge-weights often have multiple MSTs. Since HGs typically have many edges with the same length (43% of edges in the HGs constructed for the FLU8 dataset have length 5 and below), there are often many MSTs for a given HG. Since our analysis revealed that many leaves represent erroneous sequences, IgEvolution attempts to construct an MST with many low-abundance leaves with the goal to remove these leaves at the next stage. Thus, given two edges with the same weight in the HG, it prioritizes one of them over another for including in the MST. Below we describe how IgEvolution redefines edge-weights to greatly reduce the number of edges with identical weights and finds an MST in the HG with new edge-weights.

IgEvolution defines the *abundance* of an edge ($v$, $w$) in the HG as the maximum of abundances of its endpoints: *abundance*($v$, $w$)=max{*abundance*($v$), *abundance*($w$)}. It further recomputes the weight of an edge ($v$, $w$) as the sum of its original weight and the reverse abundance of this edge: *weight**($v$, $w$) = *weight*($v$,$w$) +1/*abundance*($v$,$w$). As a result, the MST algorithm prioritizes a high abundance edge over a low-abundance edge even if these edges have the same weight in the HG (ties between edges with the same weight and the same abundance are broken arbitrarily).

When the Prim algorithm decides which of two "competing" equally-weighted edges ($v$,$w$) and ($u$,$w$) to add to the growing tree, it adds an edge with higher abundance. Since selecting such edge results in a higher chance for the vertex $w$ to be classified as a low-abundance leave, it has a higher chance to be deleted at the follow-up leaf-deletion step.

We refer to an MST constructed without and with edge prioritization as *simple* and *max-leaf*, respectively (Figure A1). On average, the max-leaf (simple) MST for the FLU1-8 dataset contains 84% (75%) of leaves before the iterative leaf-removal step (Figure A2). After the leaf-removal step, the max-leaf (simple) MSTs contain 19% (45%) of low-abundance sequences on average (Figure A2).

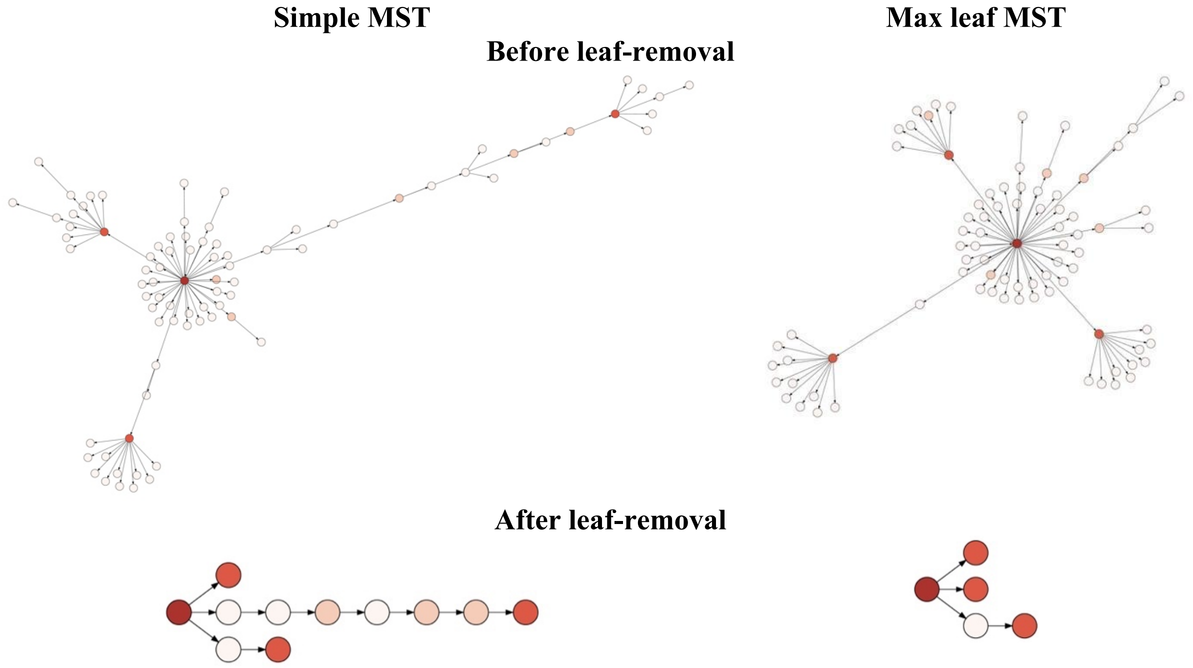


**Figure A1. Simple and max-leaf MSTs constructed for a clonal lineage from the FLU1-8 dataset.** Vertices are colored according to their abundances (from pale-low to dark-high).

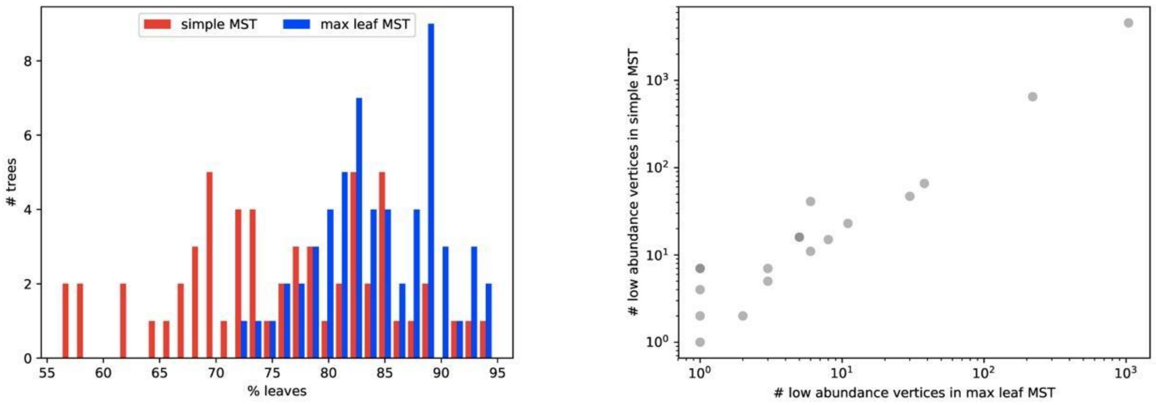


**Figure A2. Information about MSTs constructed for 58 largest clonal lineages (with at least 100 sequences) in the FLU1-8 dataset.** (Left) Distribution of the percentage of leaves in the simple MSTs (red) and the max-leaf MSTs (blue). (Right) Number of low-abundance sequences in the max-leaf ($x$-axis) and simple ($y$-axis) MSTs constructed for the same clonal lineages. Each lineage corresponds to a dot.

#### Supplemental Note: IgEvolution limitations

Although Hamming graph allows us to apply efficient algorithms (e.g., fast algorithms for Hamming graph construction), it limits our analysis to sequences with the same length. Hamming graphs work for most human, mouse, and rat datasets. In the FLU1-4 dataset, on average, 98% of sequences from the same clonal lineage have the same length. However, it becomes critical for species whose mutation process includes frequent insertions and deletions (e.g., rabbit). Thus, in cases of indels, IgEvolution tends to decompose clonal lineages into a set of smaller lineages.

In future, we plan to address this issue and modify IgEvolution in order to deal with sequences of different lengths. We also plan to extend the algorithm and reconstruct missing vertices in the clonal graph. Both these modifications involve the design of more sophisticated algorithms and we believe deserve to be described as a separate paper.

#### Supplemental Note: IgEvolution parameters

**Setting a threshold for detecting low-abundance receptor sequences.** In Shlemov et al., 2017, we showed that the most reliable absolute threshold for filtering low-abundance sequences is $minsize_{abs} = 5$. Thus, as a first step, IgEvolution iteratively removes all leaves with multiplicity at most 5. However, this procedure is not able to deal with artificial sequences resulted from PCR of highly abundant sequences.

As a second step, IgEvolution removes a leaf if the ratio of its multiplicity to the multiplicity of its paternal sequence is below $minsize_{rel}$ (by default, $minsize_{rel} = 0.05$). Figure A3 shows that the distribution of $minsize_{rel}$ values are bimodal and the default value 0.05 separates the left and the right modes. The second step is also performed in an iterative manner.

**Finding low-multiplicity leaves in SHM graphs.** To find the threshold identifying low-multiplicity leaves in SHM graphs, we computed the distribution of multiplicities of all leaves in SHM graphs for FLU1, FLU2, and INTESTINAL datasets (Figure A3). Threshold equal to 5 allows us to remove 83% of leaves.

**Finding highly mutable positions.** To identify positions with high mutability, we collected mutability values across all positions corresponding to CDR1, CDR2, and CDR3 computed according to the IMGT notation (Figure A3). The median value ($mut_{med}$) of the computed distribution is 0.031. Thus, we consider a position is highly-mutable if it is mutability exceeds $mut_{med}$(i.e., is higher that average mutability of CDR positions).

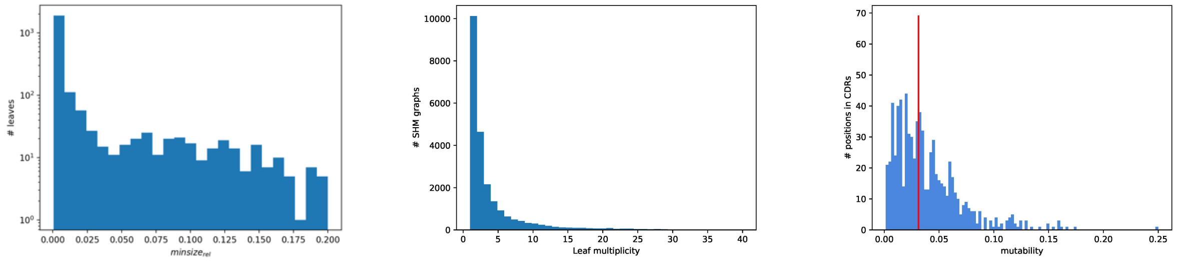


**Figure A3. IgEvolution parameters.** (Left) Distribution of *minsize*$_{rel}$ values in the FLU1-4 dataset. Only values below 0.2 are shown. 25% of *minsize*$_{rel}$ exceed 0.2. (Middle) Distribution of leaf multiplicities in the SHM graphs computed for FLU1, FLU2, and INTESTINAL datasets. (Right) Distribution of mutabilities of positions from CDRs in 78 large clonal graphs computed across FLU1, FLU2, and INTESTINAL datasets. Red line shows the median of the distribution that is equal to 0.031.

#### Supplemental Note: Performance of repertoire construction tools on clonally expanded datasets

In Shlemov et al., 2017, we compared the performance of three repertoire construction tools (IgReC, pRESTO, and MiXCR) on various types of immunosequencing datasets. While the performance of IgReC and MiXCR was higher on datasets with high VDJ diversity (e.g., PBMC of healthy individuals), pRESTO demonstrated better results on datasets with large clonal lineages (e.g., specific immune repertoires). Thus, we tested the performance of pRESTO on *FluGraph* shown in Figure 2. pRESTO strategy merges identical reads and uses a fixed threshold (*minsize*) for filtering sequences with low abundance. Figure A4 shows that any choice of the *minsize* threshold remove vertices connecting the centers of the star subgraphs. Figure A4 also shows that *minsize* equals to 5 or 10 does not remove vertices surrounding the center of the star subgraphs that might represent PCR artifacts.

#### Supplemental Note: Types of clonal graphs

Figure A5 shows the largest clonal graphs from the FLU1-6 and FLU1-8 datasets. In contrast to the largest clonal graph from the FLU1-4 dataset illustrated in the main text, these graphs contain cycles. Figure A5 shows that most cycles (shown in red) the relatively short (2–4 vertices) and do not break tree-like structure of the graph. We also see that some vertices (shown in purple) have more than one incoming edge. We assume that these vertices emerged as a result of convergent evolution in independent branches of the same clonal graph.

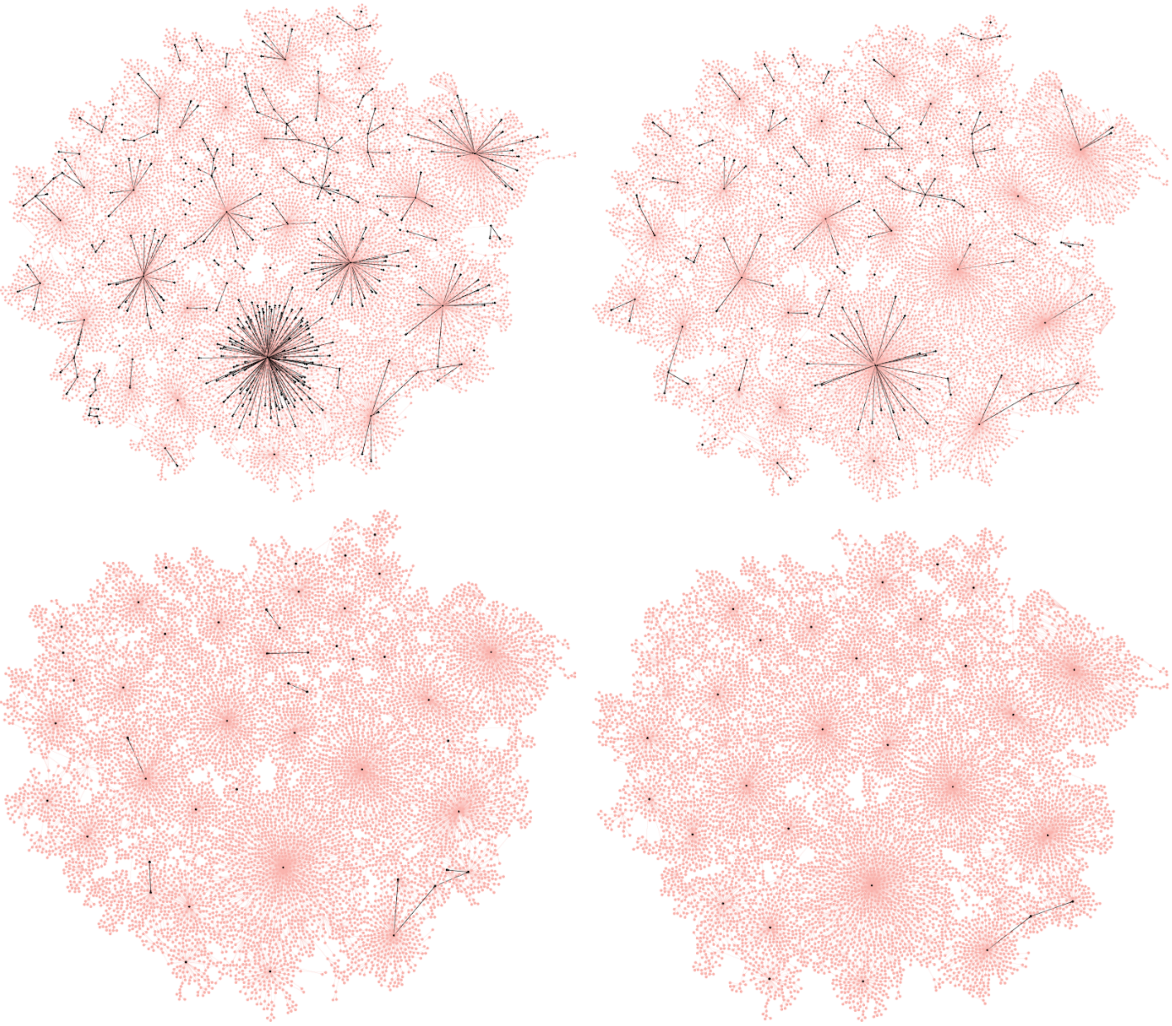

**Figure A4. Performance of pRESTO on *FluGraph* with various *minsize* thresholds: 5 (upper left), 10 (upper right), 50 (lower left), and 100 (lower right).** Black vertices correspond to sequences survived after filtering low-abundance sequences. Black edges connect black vertices.

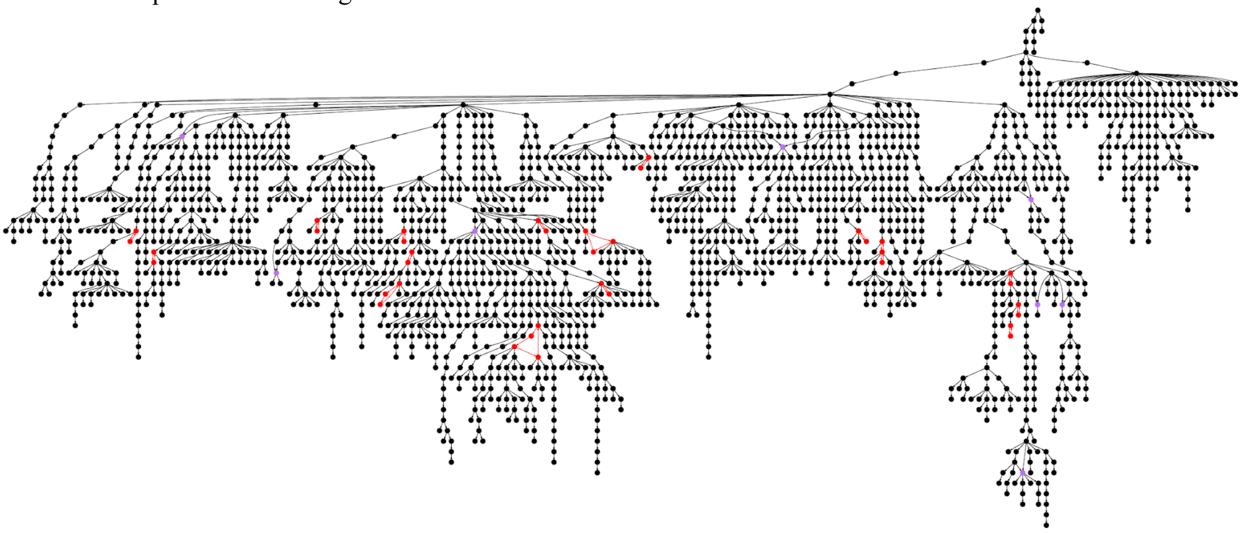

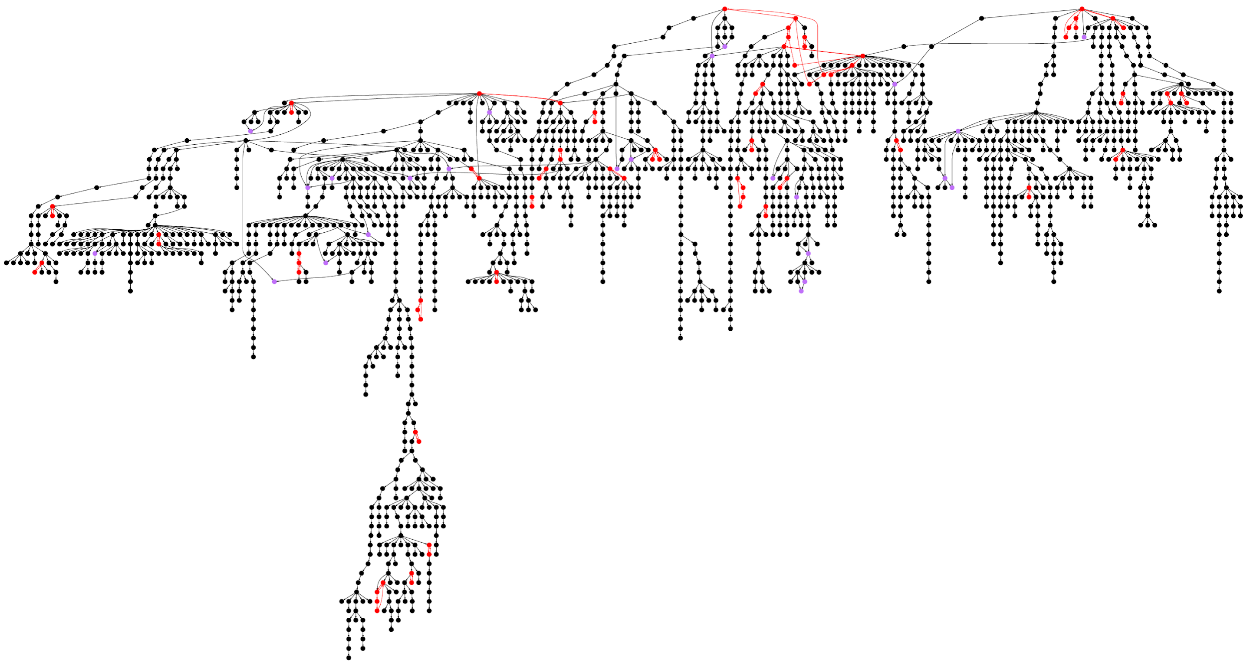

**Figure A5. The largest clonal graphs in the FLU1-6 (top) and FLU1-8 (bottom) datasets contain cycles.** Vertices corresponding to strongly connected components are colored in red. A vertex with at least two incoming edges is colored in purple (if it does not belong to a strongly connected component).

##### Supplemental Note: Test immunosequencing datasets

The FLU1 datasets are described in Tables A1, A2, A3, and A4. The FLU2 datasets are described in Table A5. The INTESTINAL datasets are described in Table A6.

| Run ID | Cell type | Time point |
|---|---|---|
| SRR3620047 | HA-positive antibody secreting cells | D7 |
| SRR3620069 | HA-positive activated B cells | D7 |
| SRR3620102 | HA-positive activated B cells | D14 |
| SRR3620028 | HA-positive memory B cells | D28 |
| SRR3620036 | naive B cells | D0 |

**Table A1. Accession numbers of the FLU1-4 dataset, project PRJNA324093, flu vaccination study.**

| Run ID | Cell type | Time point |
|---|---|---|
| SRR3620037 | HA-positive antibody secreting cells | D7 |
| SRR3620039 | HA-positive activated B cells | D7 |
| SRR3620042 | HA-positive activated B cells | D14 |
| SRR3620046 | HA-positive memory B cells | D28 |
| SRR3620035 | naive B cells | D0 |

**Table A2. Accession numbers of the FLU1-5 dataset, project PRJNA324093, flu vaccination study.**

| Run ID | Cell type | Time point |
|---|---|---|
| SRR3620055 | HA-positive antibody secreting cells | D7 |
| SRR3620057 | HA-positive activated B cells | D7 |
| SRR3620061 | HA-positive activated B cells | D14 |
| SRR3620065 | HA-positive memory B cells | D28 |
| SRR3620054 | naive B cells | D0 |

**Table A3. Accession numbers of the FLU1-6 dataset, project PRJNA324093, flu vaccination study.**

| Run ID | Cell type | Time point |
|---|---|---|
| SRR3620074 | HA-positive antibody secreting cells | D7 |
| SRR3620076 | HA-positive activated B cells | D7 |
| SRR3620079 | HA-positive activated B cell | D14 |
| SRR3620084 | HA-positive memory B cells | D28 |
| SRR3620073 | naive B cell | D0 |

**Table A4. Accession numbers of the FLU1-8 dataset, project PRJNA324093, flu vaccination study.**

| Run ID | Donor ID |
|---|---|
| SRR8377659 | Donor 1 |
| SRR8377660 | Donor 3 |
| SRR8377663 | Donor 4 |
| SRR8377656 | Donor 5 |

**Table A5. Accession numbers of the FLU2 datasets, project PRJNA512111, flu vaccination study.**

| Run ID | Cell line | Isotype | Tissue | Run ID | Cell line | Isotype | Tissue |
|---|---|---|---|---|---|---|---|
| **Donor 1** | | | | **Donor 3** | | | |
| SRR5063099 | memory B cells | IgA | ileum mucosa | SRR5063086 | memory B cells | IgA | colon mucosa |
| SRR5063100 | plasma B cells | IgA | colon mucosa | SRR5063096 | memory B cells | IgA | ileum mucosa |
| SRR5063104 | memory B cells | IgA | colon mucosa | SRR5063101 | plasma B cells | IgA | colon mucosa |
| SRR5063107 | plasma B cells | IgA | ileum mucosa | SRR5063106 | plasma B cells | IgA | ileum mucosa |
| SRR5063089 | plasma B cells | IgM | ileum mucosa | SRR5063087 | plasma B cells | IgM | colon mucosa |
| SRR5063090 | plasma B cells | IgM | colon mucosa | SRR5063094 | memory B cells | IgM | colon mucosa |
| SRR5063105 | memory B cells | IgM | ileum mucosa | SRR5063102 | memory B cells | IgM | ileum mucosa |
| SRR5063114 | memory B cells | IgM | colon mucosa | SRR5063103 | plasma B cells | IgM | ileum mucosa |
| **Donor 2** | | | | **Donor 4** | | | |
| SRR5063082 | plasma B cells | IgA | ileum mucosa | SRR5063088 | plasma B cells | IgA | colon mucosa |

| | | | | | | | |
|---|---|---|---|---|---|---|---|
| SRR5063083 | plasma B cells | IgA | colon mucosa | SRR5063091 | plasma B cells | IgA | ileum mucosa |
| SRR5063093 | memory B cells | IgA | colon mucosa | SRR5063109 | memory B cells | IgA | ileum mucosa |
| SRR5063115 | memory B cells | IgA | ileum mucosa | SRR5063116 | memory B cells | IgA | colon mucosa |
| SRR5063095 | plasma B cells | IgM | ileum mucosa | SRR5063085 | memory B cells | IgM | colon mucosa |
| SRR5063098 | memory B cells | IgM | colon mucosa | SRR5063108 | plasma B cells | IgM | colon mucosa |
| SRR5063111 | memory B cells | IgM | ileum mucosa | SRR5063110 | plasma B cells | IgM | ileum mucosa |
| SRR5063113 | plasma B cells | IgM | colon mucosa | SRR5063112 | memory B cells | IgM | ileum mucosa |

**Table A6. Accession numbers of the INTESTINAL datasets, project PRJNA355402, intestinal antibody repertoire study.**

##### Supplemental Note: Clonal analysis of rat antibody repertoires

We applied IgEvolution to PBMC samples taken from 10 rats after vaccination with DNP or HuD immunogens (project PRJNA386462, VanDuijn et al., 2017). We will refer to these datasets as RAT (Table A7). In total, IgEvolution computed 76 large clonal graphs (at least 50 vertices). Figure A6 shows that the mutability of CDRs and FRs in the RAT datasets looks similar to the human datasets. This suggests that observations about the function of CDRs and FRs described in text main text can be also applied for rat antibodies. In future, we plan to apply IgEvolution to other non-human species and perform comparative analysis of clonal development of antibody repertoires in vertebrate species.

| Run ID | Individual | Immunogen | Run ID | Individual | Immunogen |
|---|---|---|---|---|---|
| SRR5534359 | 1 | DNP | SRR5534364 | 6 | HuD |
| SRR5534360 | 2 | DNP | SRR5534365 | 7 | HuD |
| SRR5534361 | 3 | DNP | SRR5534366 | 8 | HuD |
| SRR5534362 | 4 | DNP | SRR5534367 | 9 | HuD |
| SRR5534363 | 5 | DNP | SRR5534368 | 10 | HuD |

**Table A7. Accession numbers of the RAT datasets, project PRJNA386462, project PRJNA386462, DNP and HuD vaccination of rats.**

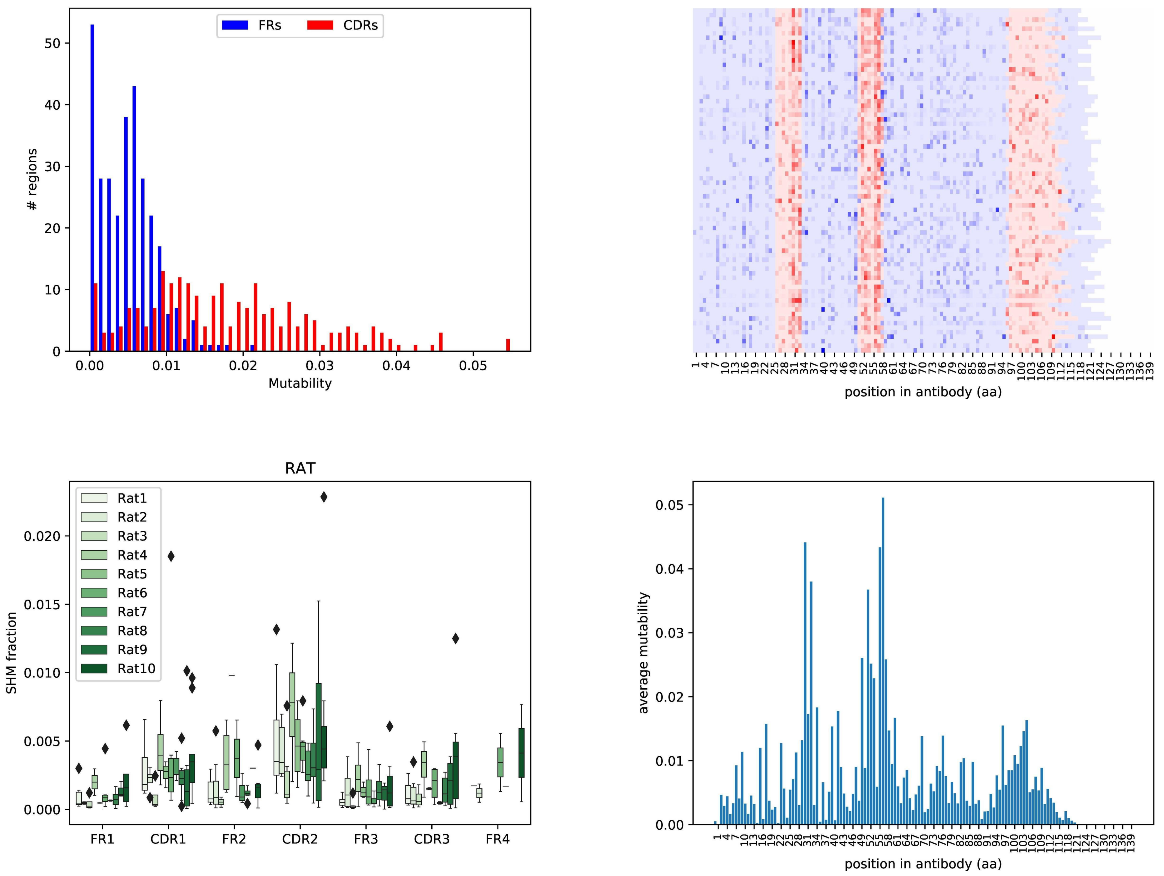


**Figure A6. Mutability of large clonal graphs computed across all RAT datasets.** All statistics are computed using 76 large clonal graphs from the RAT datasets. (Left) Distribution of mutability values of CDRs and FRs. (Right) Mutability maps. The detailed description of the plots is provided the captions to Figures 4 and 5.

**Supplemental Note: Finding binding and contact sites in antibody-antigen complexes**

We downloaded 953 antibody-antigen complexes from the SAbDab database (Dunbar et al., 2014) in PDB format. To identify positions in heavy chains corresponding to antigen binding sites and light chain contacts, we applied BioPython package for reading PDB files and extracted intermolecular residues located at the distance <5A. 745 out of 953 structures have complete PDB files and were processed by BioPython without errors.

**Supplemental Note: Analyzing highly mutable positions in typical FR1, FR2, and CDR2**

**Analyzing highly mutable positions in FR1.** Surprisingly, although FR1 contains few binding and contact sites (< 4% of the 3D structures), positions 6, 12, 23, and 24 in FR1 are highly mutable in more than $perc_{med}$ (9–27%) of clonal graphs. The last two positions (23 and 24) are also characterized by many convergent SHMs.

**Analyzing highly mutable positions in FR2.** Positions corresponding to the frequent contact sites in FR2 (positions 4–15) have low mutabilities in clonal graphs: on average, these positions are highly-mutable in only 2% of the clonal graphs. On the other hand, positions 1, 2, 17 (likely presenting extensions of binding sites located in CDR1 and CDR2) represent highly mutable positions in more than $perc_{med}$ (8–28%) of clonal graphs. We thus assume that an amino acid at a contact site undergoes a strong selection against changing this amino acid.

**Analyzing highly mutable positions in CDR2.** All eight positions in CDR2 represent binding sites in many 3D structures (18–53%). Five of them (3, 4, 6, 7, 8) are also highly mutable in more than $perc_{med}$

(8–27%) of clonal graphs. These five positions are characterized by many distinct SHMs and few convergent SHMs. We assume that such features are typical for binding sites.

**Supplemental Note: Alleles of IGHV3-11 and IGHV4-39 shape immunoglobulin response**

Clonal usage of V genes reveals the differences between repertoire with the same specificity (Figure 7). We showed that IGHV1-69 that is known for having high affinity to hemagglutinin is utilized by donors 4 and 8, but missing in donors 5 and 6. In the main text, we showed that the differences in usage are explained by genomic variations. Figure 7 also shows that other V genes are also used differently in donors from the FLU1 datasets: IGHV3-11 is used by donors 4 and 5 but missing in donors 6 and 8; IGHV4-39 is used by donors 4, 5, and 6, but missing in the donor 8. We assume that these differences can also be caused by genomic variations of the genes. Thus, we applied the same analysis to IGHV3-11 and IGHV4-39 as we did for IGHV1-69. Figure A7 shows that differences in usage of IGHV3-11 and IGHV4-39 can be explained by genomic variations.

**Clonal analysis of IGHV3-11.** For IGHV3-11, we selected five positions corresponding to genomic variations: 55, 57, 58, 59, and 72. Donors 4 and 5 have the same amino acids that these positions (G–T–I–Y–R) corresponding to a homozygous allele of IGHV3-11. We refer to this allele as dominant. Donor 8 has different amino acids at the selected positions (S–Y–T–N–K) corresponding to another homozygous allele of IGHV3-11. Donor 6 has a heterozygous allele of IGHV3-11 presenting a mixture of two these homozygous alleles (G/S–T/Y–I/T–Y/N–R/K). Figure A8 shows amino acid content at the selected positions in non-trivial clonal graphs derived from IGHV3-11 in the FLU1 and FLU2 datasets (there are no non-trivial clonal graphs derived from IGHV3-11 in the INTESTINAL datasets). Figure A8 shows that only position 72 preserves the dominant amino acid (Arg). Thus, we assume that position 72 differentiates usage of IGHV3-11 in the flu-specific immune response.

**Clonal analysis of IGHV4-39.** For IGHV4-39, we also selected five positions corresponding to genomic variations: 31, 33, 55, 78, and 83. Donors 4, 5, 6 have the same amino acids that these positions (S–S–Y–N–K) corresponding to a homozygous allele of IGHV4-39. We refer to this allele as dominant. Donor 8 possesses a heterozygous allele of the gene representing a mixture of the dominant allele and an unknown allele (S/R–S/T–Y/F–N/K–K/R). Figure A9 shows amino acid content at the selected positions in non-trivial clonal graphs derived from IGHV4-39 in the FLU1, FLU2, and INTESTINAL datasets. Figure A9 shows that the dominant amino acids are preserved at positions 55 (Tyr) and 78 (Asn). Thus, we assume that positions 55 and 78 differentiates usage of IGHV4-39 in antigen specific immune response.

**IGHV3-11**

| AA position | 55 | 57 | 58 | 59 | 72 |
|---|---|---|---|---|---|
| Donor 4 | G | T | I | Y | R |
| Donor 5 | G | T | I | Y | R |
| Donor 6 | G / S | T / Y | I / T | Y / N | R / K |
| Donor 8 | S | Y | T | N | K |

```
IGHV3-11*01  QVQLVESGGGLVKPGGSLRLSCAASGFTFSDYYMSWIRQAPGKGLEWVSYISSSGSTIYYADSVKGRFTISRDNAKNSLYLQMNSLRAEDTAVYYCAR
IGHV3-11*02  QVQLLESGGGLVKPGGSLRLSCAASGFTFSDYYMSWIRQAPGKGLEWVSYISSSSSYTNYADSVKGRFTISRDNAKNSLYLQMNSLRAEDTAVYYCAR
IGHV3-11*03  QVQLVESGGGLVKPGGSLRLSCAASGFTFSDYYMSWIRQAPGKGLEWVSYISSSGSTIYYADSVKGRFTISRDNAKNSLYLQMNSLRAEDTAVYYCAR
IGHV3-11*04  QVQLVESGGGLVKPGGSLRLSCAASGFTFSDYYMSWIRQAPGKGLEWVSYISSSSSYTNYADSVKGRFTISRDNAKNSLYLQMNSLRAEDTAVYYCAR
IGHV3-11*05  QVQLVESGGGLVKPGGSLRLSCAASGFTFSDYYMSWIRQAPGKGLEWVSYISSSSSYTNYADSVKGRFTISRDNAKNSLYLQMNSLRAEDTAVYYCAR
```

**IGHV4-39**

| AA position | 31 | 33 | 55 | 78 | 83 |
|---|---|---|---|---|---|

| | | | | | |
|---|---|---|---|---|---|
| **Donor 4** | **S** | **S** | **Y** | **N** | **K** |
| **Donor 5** | **S** | **S** | **Y** | **N** | **K** |
| **Donor 6** | **S** | **S** | **Y** | **N** | **K** |
| **Donor 8** | **S / R** | **S / T** | **Y / F** | **N / K** | **K / R** |

**Figure A7. Germline mutations in IGHV3-11 (upper) and IGHV4-39 (lower) in the FLU1 datasets.** (Lower) 6 out of 7 positions detected in naive datasets as genomic variations change the amino acid sequence of IGHV3-11 resulting in 5 pairs of dominant and recessive amino acids: 55: G/S, 57: T/Y, 58: I/T, 59: Y/N, and 72: R/K. 4 out of 5 positions with genomic variations are confirmed by known alleles of IGHV3-11. (Upper) 5 out of 9 positions detected in naive datasets as genomic variations change the amino acid sequence of IGHV4-39 resulting in 5 pairs of dominant and recessive amino acids: 31: S/R, 33: S/T, 55: Y/F, 78: N/K, and 83: K/R. None of the positions is confirmed by known alleles of IGHV4-39.

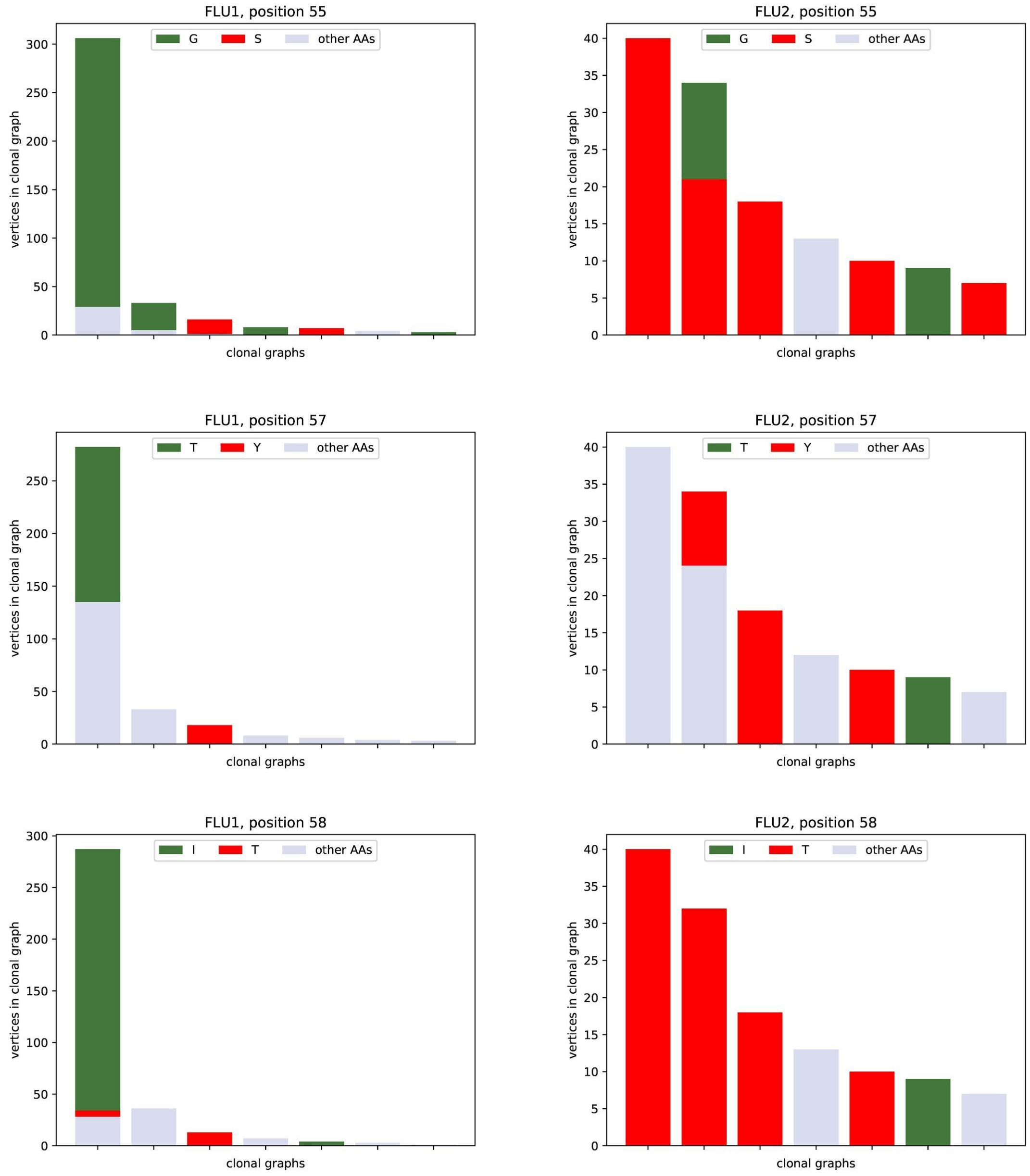

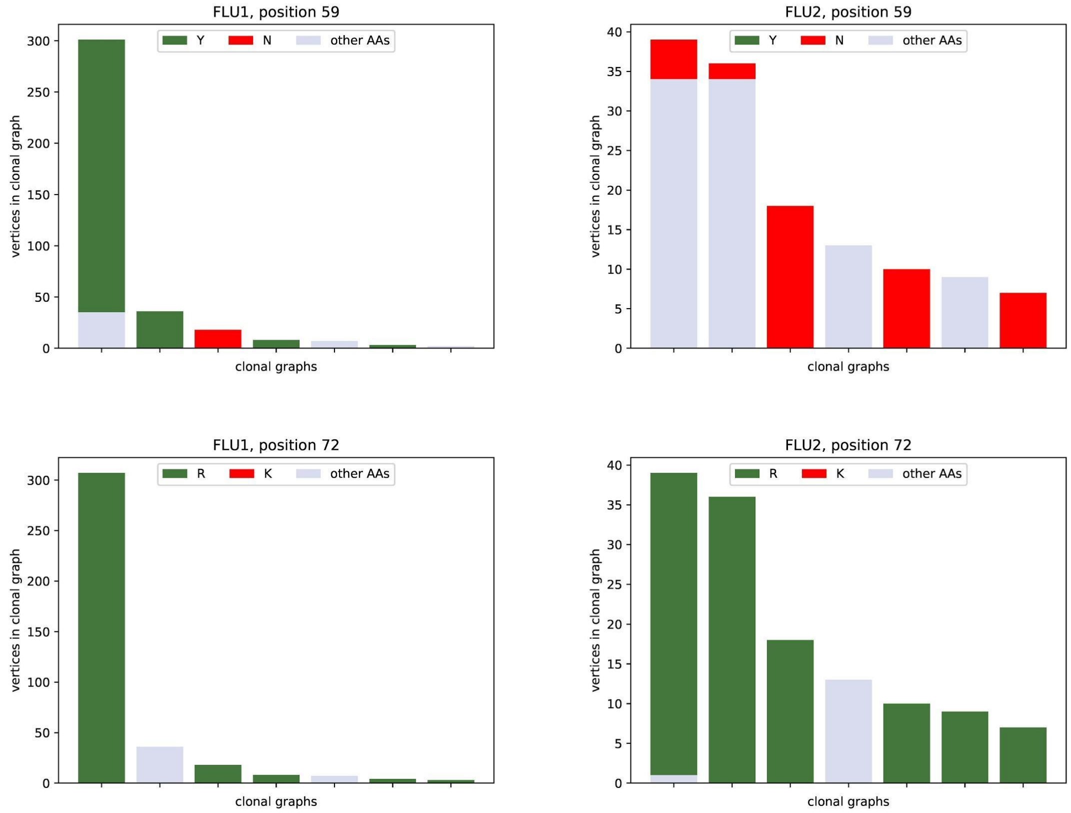


**Figure A8. Amino acid content corresponding to positions 55, 57, 58, 59, and 72 in clonal graphs derived from IGHV3-11 in the FLU1 (left) and FLU2 (right) datasets, respectively.**

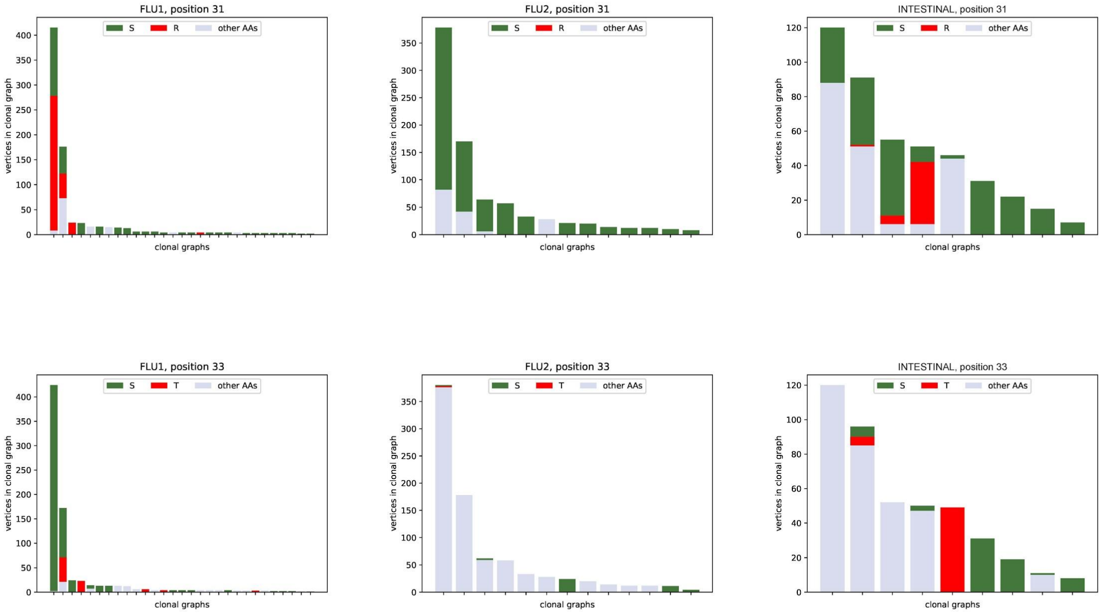

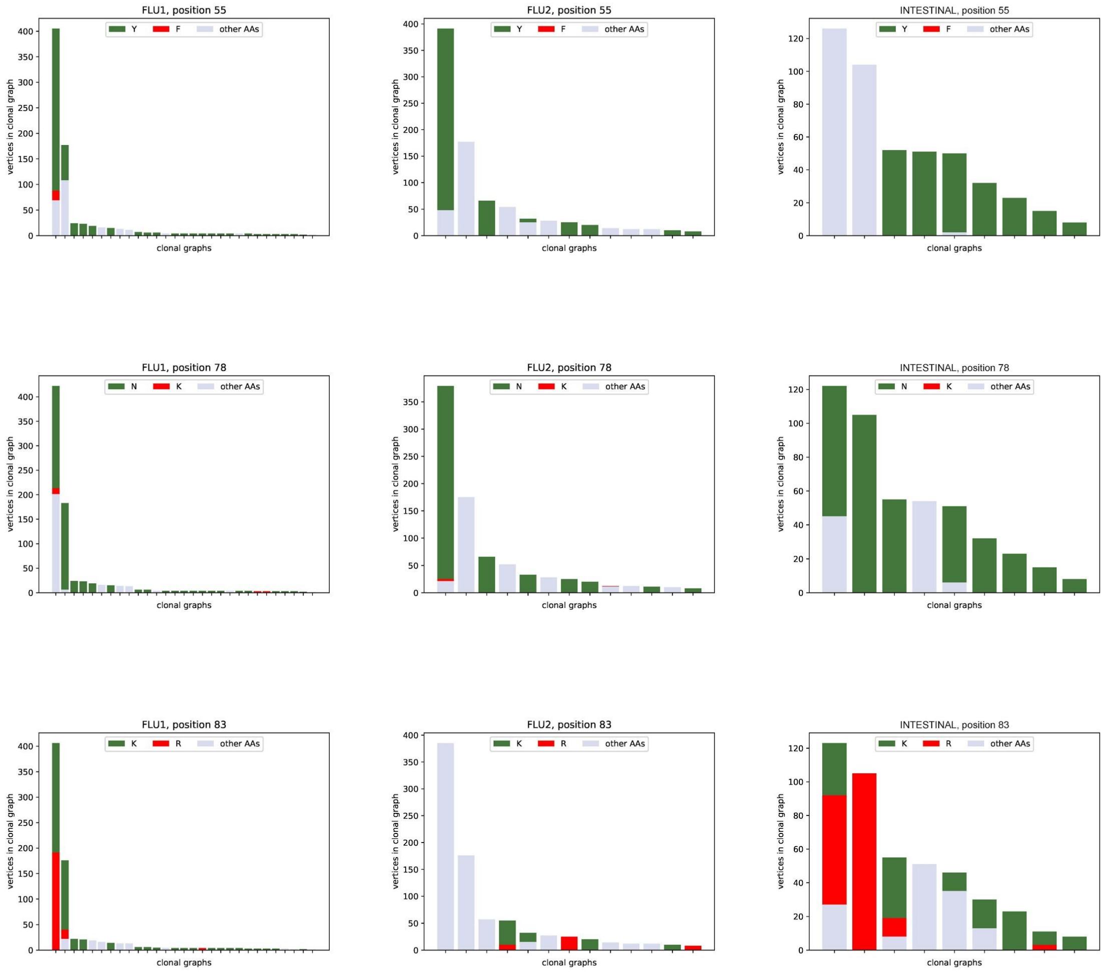


**Figure A9. Amino acid content corresponding to positions 31, 33, 55, 78, and 83 in clonal graphs derived from IGHV4-39 in the FLU1 (left), FLU2 (middle), and INTESTINAL (right) datasets, respectively.**
